# Perilipin2 down-regulation in β cells impairs insulin secretion under nutritional stress and damages mitochondria

**DOI:** 10.1101/2020.10.01.322974

**Authors:** Akansha Mishra, Siming Liu, Joseph Promes, Mikako Harata, William Sivitz, Brian Fink, Gourav Bhardwaj, Brian T. O’Neill, Chen Kang, Rajan Sah, Stefan Strack, Samuel Stephens, Timothy King, Laura Jackson, Andrew S Greenberg, Frederick Anokye-Danso, Rexford S Ahima, James Ankrum, Yumi Imai

**Affiliations:** Department of Internal Medicine, Carver College of Medicine, University of Iowa, Iowa City, IA; Fraternal Order of Eagles Diabetes Research Center, University of Iowa, Iowa City, IA; Iowa City Veterans Affairs Medical Center, Iowa City, IA; Department of Internal Medicine, Washington University School of medicine, St Louis, MO; Department of Neuroscience and Pharmacology, Iowa Neuroscience Institute University of Iowa, Iowa City, IA; Department of Internal Medicine, Eastern Virginia Medical School, Norfolk, VA; Obesity and Metabolism Laboratory, Jean Mayer United States Department of Agriculture Human Nutrition Research Center on Aging at Tufts University, Boston, MA; Department of Medicine, Johns Hopkins School of Medicine, Baltimore, MD; Roy J. Carver Department of Biomedical Engineering, University of Iowa, Iowa City, IA

## Abstract

Perilipin 2 (PLIN2) is the lipid droplet (LD) protein in β cells that increases under nutritional stress. Down-regulation of PLIN2 is often sufficient to reduce LD accumulation. To determine whether PLIN2 positively or negatively affects β cell function under nutritional stress, PLIN2 was down-regulated in mouse β cells, INS1 cells, and human islet cells. β cell specific deletion of PLIN2 in mice on a high fat diet reduced glucose-stimulated insulin secretion (GSIS) in vivo and in vitro. Down-regulation of PLIN2 in INS1 cells blunted GSIS after 24 h incubation with 0.2 mM palmitic acids. Down-regulation of PLIN2 in human pseudoislets cultured at 5.6 mM glucose impaired both phases of GSIS, indicating that PLIN2 is critical for GSIS. Down-regulation of PLIN2 decreased specific OXPHOS proteins in all three models and reduced oxygen consumption rates in INS1 cells and mouse islets. Moreover, we found that PLIN2 deficient INS1 cells increased the distribution of a fluorescent oleic acid analog to mitochondria and showed signs of mitochondrial stress as indicated by susceptibility to fragmentation and alterations of acyl-carnitines and glucose metabolites. Collectively, PLIN2 in β cells have an important role in preserving insulin secretion, β cell metabolism and mitochondrial function under nutritional stress.

## Introduction

It is widely accepted that the progressive loss of functional β cell mass is a key pathology of type 2 diabetes (T2D) (1, 2). Nutritional stress due to increased influx of glucose and lipids is considered to trigger and accelerate β cell dysfunction in T2D by activating stress pathways including endoplasmic reticulum (ER) stress, oxidative stress, mitochondrial dysfunction, and inflammation (3, 4). Thus, it is important to understand how β cells respond to increase in nutritional influx. One consequence of nutritional overload is the increased production of lipids including triglycerides (TG) that are stored in lipid droplets (LDs) (5). While the accumulation of LDs is closely tied to tissue dysfunction in non-alcoholic fatty liver disease, insulin resistant skeletal muscle, atherosclerosis, and diabetic cardiomyopathy, a role of LD under nutritional stress in β cell dysfunction has been underappreciated due to difficulty demonstrating LDs in mouse β cells (3, 6, 7). Importantly, LDs are easily identifiable in adult human β cells (6, 8, 9). Active formation of LDs in human β cells was shown not only *in vitro* (9) but also *in vivo* utilizing human islet transplantation in immune deficient mice on high fat diets (HFD) (10). However, it is currently unknown whether LDs in β cells are merely a marker of nutritional overload or do they play a role in β cell function under physiology and nutritional stress.

The perilipin (PLIN) family of proteins reside on the surface of LDs and regulate intracellular lipid metabolism by controlling the LD formation and mobilization, as well as the interaction with other organelles (11). Among PLINs, PLIN2 is the most abundant PLIN in many non-adipocytes including β cells (4). We previously demonstrated that PLIN2 is increased in parallel to TG accumulation in islets of mice fed HFD, in *ob/ob* mice, and in human islets after fatty acid (FA) loading (12); findings also reported by others (8, 13). Recently, an increase in immunostaining of PLIN2 was reported in human T2D islets implicating PLIN2 as a possible key PLIN in human T2D (14). PLIN2’s role is not limited to passively affecting lipid accumulation. Rather, PLIN2 actively regulates LD accumulation in a β cell model: the down-regulation of PLIN2 by antisense oligonucleotides (ASO) decreased while overexpression of PLIN2 increased FA incorporation into TG in MIN6 cells (9, 12). Thus, the down-regulation of PLIN2 should allow us to address whether LDs play a positive or negative role during the development of β cell failure under nutritional stress. Previously, whole body PLIN2 knockout (KO) mice (exon 2-3 deletion, Δ2-3) carrying Akita insulin mutation demonstrated better preservation of β cell mass and reduced ER stress suggesting that PLIN2 may protect β cells under extreme ER stress (13). Although ER stress likely has an important role in β cell demise in T2D (15), it remains unknown whether PLIN2 in β cells accelerates or prevents β cell dysfunction under nutritional stress. Considering that down-regulation of PLIN2 in MIN6 cells under regular culture conditions impaired acute augmentation of insulin secretion by palmitic acids (PA) (12), it is important to address whether the reduction of PLIN2 improves β cell function under nutritional stress.

Here, we demonstrated that β cell specific deletion of PLIN2 (exon 5 deletion, Δ5) caused the impairment of insulin secretion both in vivo and in vitro in mice on HFDs. Impairment in insulin secretion was also seen in INS1 cells and human pseudoislets after the down-regulation of PLIN2 indicating that PLIN2 supports insulin secretion especially under nutritional overload. Increased targeting of FA to mitochondria with associated mitochondrial dysfunction were implicated as the basis for β cell dysfunction after PLIN2 down-regulation.

## Results

### Insulin secretion is impaired in β cell specific PLIN2 knockout mice on high fat diet

To address whether the increase of PLIN2 seen in β cells under overnutrition is protective or detrimental to β cells, we placed β cell specific PLIN2 KO mice (βKO) on HFDs, a model that previously resulted in increased TG and PLIN2 levels in islets (12). As expected, islets from βKO mice showed significantly reduced PLIN2 mRNA and protein levels (Fig. 1a, b). Body weight (BW) and glucose tolerance did not differ between male WT and βKO mice on regular rodent chow (Supplementary Fig 1a, b). While 3 months of HF feeding increased BW (Fig. 1c vs Supplementary Fig 1c) and blood glucose (Fig. 1d vs Supplementary Fig 1b, time 0), PLIN2 deficiency in β cells did not confer better glucose tolerance (Fig. 1d) or reduced fasting glucose (Fig. 1e). There was also no significant change in insulin tolerance between WT and βKO mice either (Fig. 1f). However, the acute rise of serum insulin in response to glucose was blunted in βKO mice compared with WT littermates (Fig. 1g, h). Female βKO mice fed HF also had reduced glucose-stimulated insulin secretion (GSIS) with no differences in BW or glucose tolerance compared with control littermates (supplementary Fig. 1c-f). Thus, the reduction of PLIN2 in mouse β cells blunts GSIS in vivo, but likely to an extent not severe enough to cause overt hyperglycemia in βKO mice fed HF.

**Figure 1.**
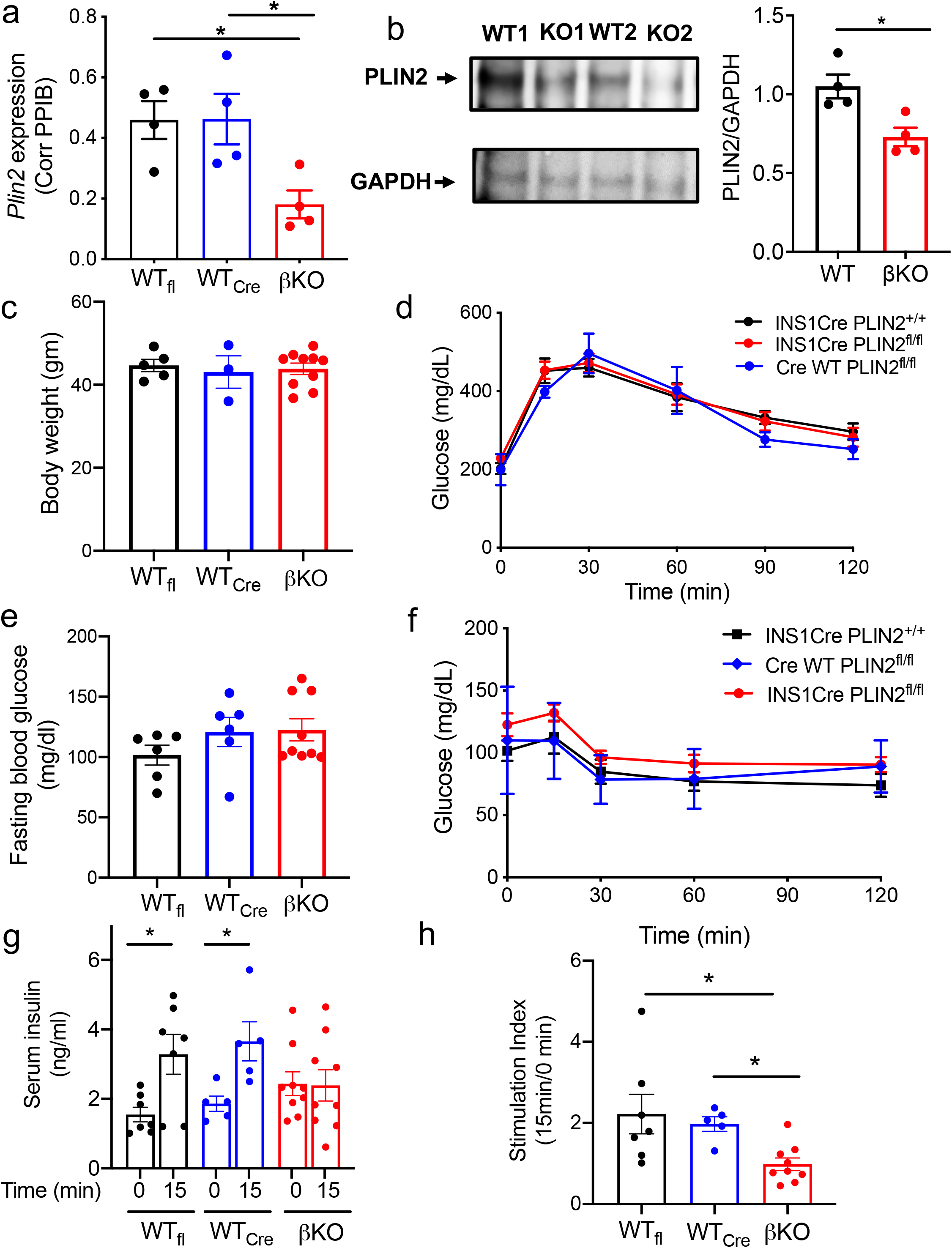
β cell specific deletion of PLIN2 impairs insulin secretion from mice on high fat diets. (a) Expression of *Plin2* in β cell specific PLIN2 KO islets compared with WT littermates determined by qPCR using *Ppib* as an internal control. WT_fl_; INS1Cre PLIN2^+/+^, WT_cre_; CreWT PLIN2^fl/fl^, βKO; INS1Cre PLIN2^fl/f^. n= 4. (b) A representative Western blot and densitometry data of PLIN2 corrected for GAPDH. (c) Body weight (BW), (d) 1.0 mg/gm BW i.p. GTT, (e) blood glucose after overnight fasting, and (f) 0.75 mU/gm BW insulin i.p. ITT on 6 month-old male mice after high fat diets for 3 months. (c, d) n= 5 for WT_fl_, 3 for WT_cre_, and 10 for βKO. (e, f) n= 6 for WT_fl_, 2 for WT_cre_, and 9 for βKO. (g) Serum insulin in response to 1.5 mg/gm BW glucose i.p. and (h) stimulation index determined as the ratio of insulin levels at 15 min to levels at 0 min for each mouse. n= 5 for WT_fl_, 7 for WT_cre_, and 9 for βKO. Data are mean ± s.e.m. for all. *; *p* <0.05 by Student’s t test.

Insulin secretion was assessed further using islets isolated from WT and βKO mice on HFDs. Here, we focused on male mice that are more susceptible to metabolic effects of HFD than females. WT islets appropriately exhibited increase in GSIS, a response augmented by acute exposure to PA (*p*<0.05 vs. 1.8 mM glucose). In contrast, βKO islets did not increase insulin secretion in response to high glucose and PA (not significant vs. 1.8 mM glucose, Fig. 2a).

**Figure 2.**
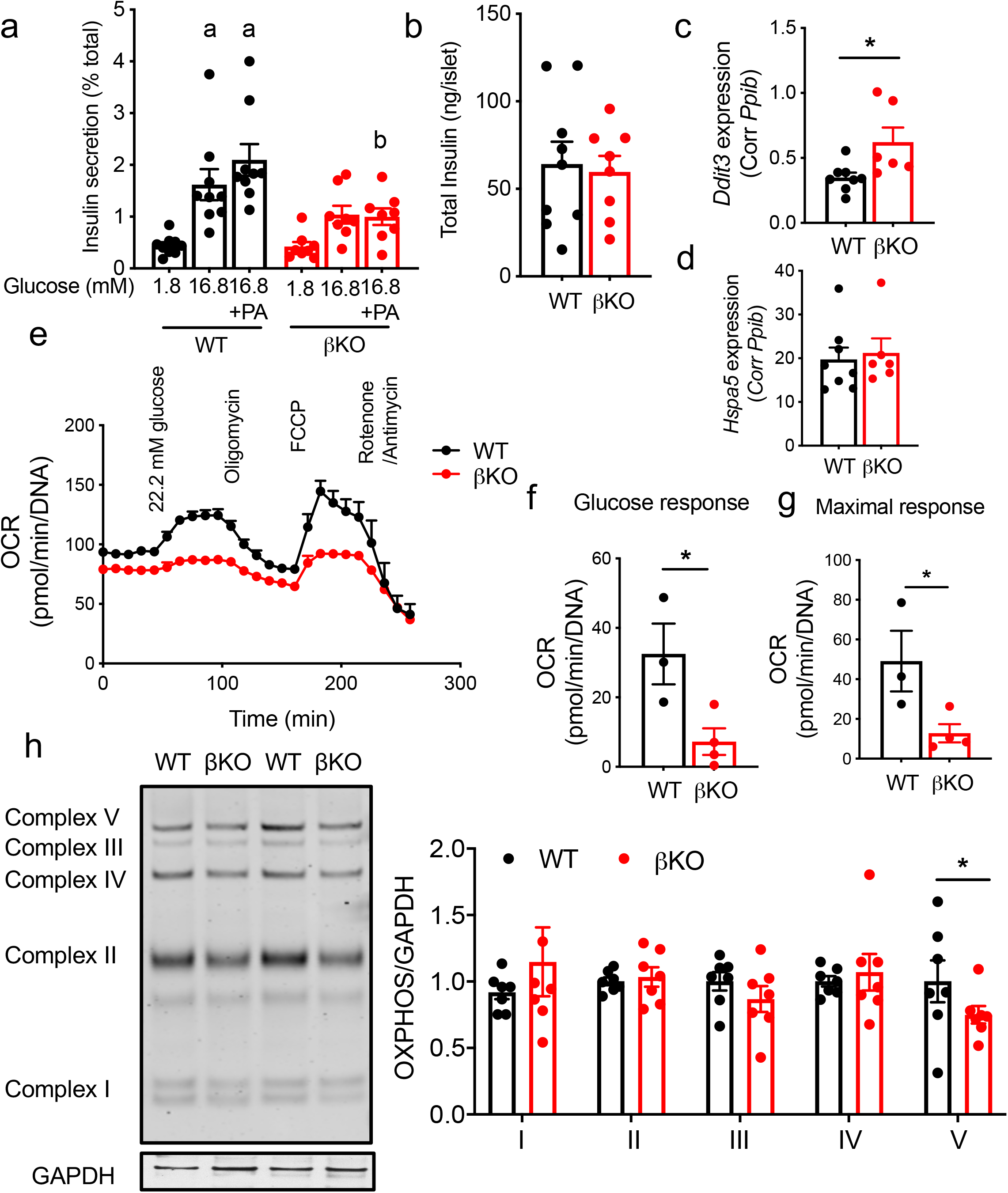
Islets from β cell specific PLIN2 knockout mice on high fat diets showed impaired insulin secretion and impaired mitochondrial function. (a) Insulin secretion measured by static incubation at indicated concentrations of glucose ± 0.5 mM PA for 1 h corrected for total insulin contents in islets of HF fed WT and β cell specific PLIN2 KO mice (βKO). (b) Total insulin contents per islet for (a). n= 9 for WT (3 WT_fl_ and 6 WT_cre_) and 8 for βKO. a; *p*< 0.05 vs. WT 1.8 mM glucose, b; *p*<0.05 vs. WT> 16.8 mM glucose + PA by One-way ANOVA with Sidak’s multi comparison test. (c, d) Expression of *Ddit3* (c) and *Hspa5* (d) in βKO and WT littermates determined by qPCR using *Ppib* as an internal control. n= 8 for WT (5 WT_fl_ and 3 WT_cre_) and 8 for βKO. (e) Oxygen consumption rate (OCR) by Seahorse Metabolic analyzer in high fat fed WT (WT_fl_) and βKO islets corrected for DNA. (f) Glucose response and (g) maximal respiration from (e). n=4 for WT_fl_ and 3 for βKO. (h) Representative Western blot and densitometry data comparing OXPHOS complexes between WT and βKO islets from high fat fed mice. Data are expressed taking average of OXPHOS/GAPDH in WT for each complex as 1. n=6 for WT (2WT_fl_ and 4 WT_cre_) and 7 for βKO. Data are mean ± s.e.m.. *; *p* <0.05 by Student’s t test.

The blunted GSIS was not associated with change in total insulin contents (Fig. 2b) or β cell area (supplementary Fig. 2a, b) in βKO islets compared with WT islets. βKO islets showed a mild increase in *Ddit3* without a change in *Hspa5* indicating PLIN2 deficiency in β cells activates a part of ER stress response under HFD (Fig. 2c, d). The impairment of GSIS in βKO islets was associated with a reduced oxygen consumption rate (OCR) in response to glucose and maximal respiration (Fig. 2e-g). The proton leak was not altered in βKO islets compared with WT islets (supplementary Fig. 2c). βKO islets also showed a significant decrease in mitochondrial complex V protein when compared with WT islets indicating defects in the mitochondrial respiratory chain (Fig. 2h). However, there was no significant difference in mitochondrial DNA between WT and βKO islets (supplementary Fig. 2d).

### PLIN2 deficiency impaired insulin secretion in INS1 cells cultured with and without palmitic acids

Data from βKO mouse islets indicated that the reduction of PLIN2 in β cells does not confer protection against HFD. However, the very small size of LDs in mouse islets and the ability of mouse β cells to compensate for HFD feeding might limit the impact of PLIN2 deficiency (10, 16). Thus, we down-regulated PLIN2 in INS1 cells, a β cell model that shows similarity to human β cells in the formation of LDs as shown in Fig. 3a and our previous study (17). Also, since PLIN2 is the almost exclusively expressed PLIN isoform in INS1 cells that express little PLIN3, INS1 cells represent an ideal model to test the contribution of PLIN2 (4). Of note, PLIN3 is known to have functional redundancy with PLIN2 (18) and is the second most abundant PLIN in mouse islets (4). After down-regulating PLIN2 using siRNA (SiPLIN2) (Supplementary Fig. 3 a-b), LDs were reduced both in number (Supplementary Fig. 3c) and size (Fig. 3a-c) in INS1 cells confirming that PLIN2 is a critical determinant of LD mass in INS1 cells. Then, PLIN2 deficient and control INS1 cells were treated with or without 0.2 mM PA for 24 h, a dose of PA chosen to cause mild cytotoxicity (19). PA treatment increased insulin secretion at low glucose in INS1 cells treated with Scr reducing the stimulation index (SI, Fig. 3d, e). Interestingly, INS1 cells after PLIN2 down-regulation showed higher basal insulin secretion and lower SI in the absence of PA treatment (Fig. 3d, e). 24h treatment with a modest dose of PA (0.2 mM), further increased basal insulin secretion. GSIS was totally lost in SiPLIN2 treated INS1 cells compared with scramble (Scr) control wherein glucose response was maintained (Fig. 3d, e). Although there was modest increase in total insulin after the downregulation of PLIN2 (Fig. 3f), this did not fully account for the increase in basal insulin secretion. Basal insulin secretion corrected for total insulin was still increased in SiPLIN2 treated INS1 cells compared with control (Fig. 3g). Considering that the increase in basal insulin secretion is an early response to nutritional stress in β cell lines, rodent islets, and human islets (20–23), the rise of basal insulin secretion in PLIN2 deficient INS1 cells in the absence of PA implicates that PLIN2 deficiency results in nutritional stress in INS1 cells in regular growth medium. Furthermore, severe blunting of GSIS after 24h exposure to PA in SiPLIN2 INS1 cells indicates that PLIN2 is protective against nutritional load in INS1 cells. In agreement with the reduced SI in SiPLIN2 INS1 cells, the rise of [Ca^2+^]_i_ in response to glucose was reduced in SiPLIN2 INS1 cells under regular culture conditions (Fig. 3h, i, Supplementary Fig. 3d).

**Figure 3.**
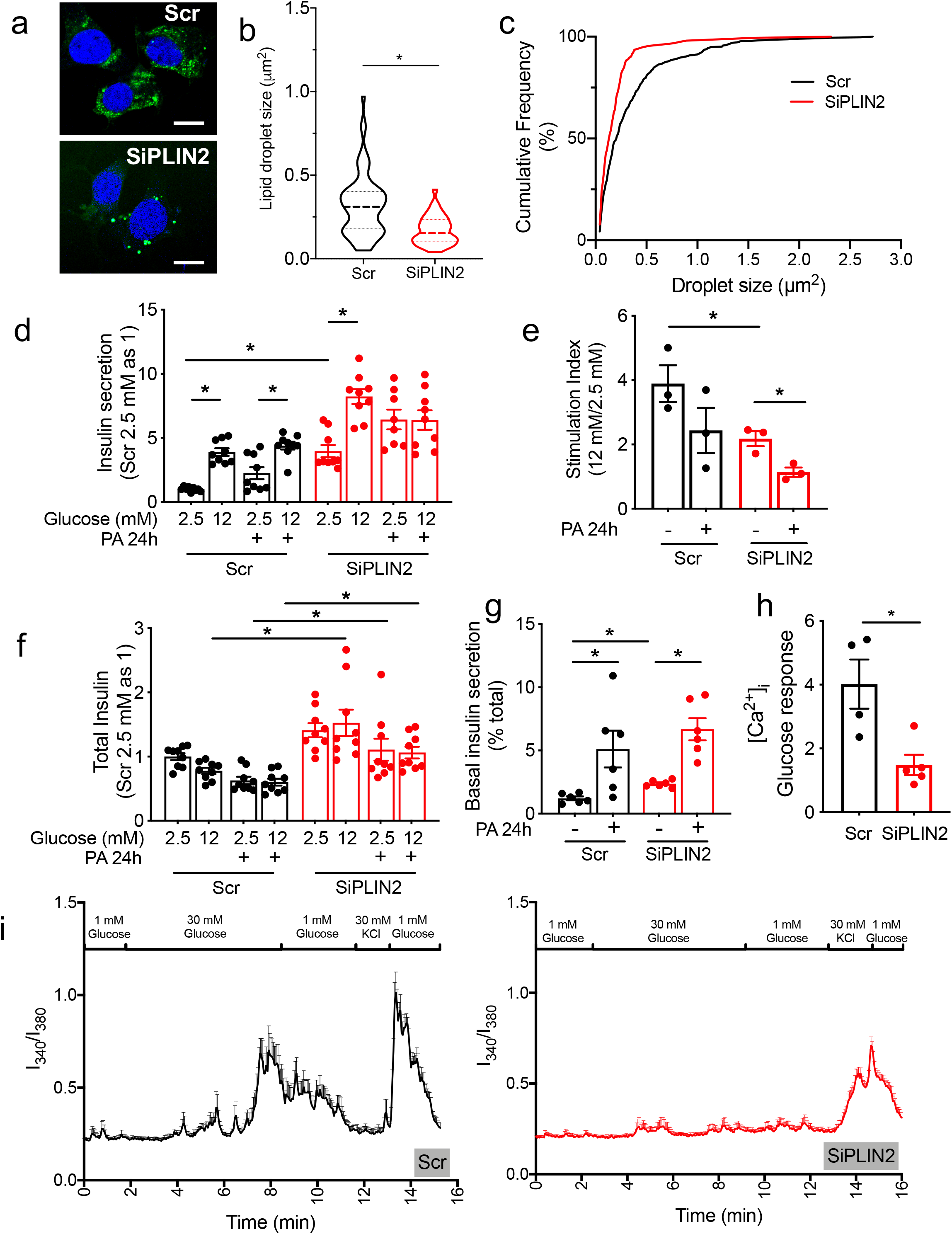
Down-regulation of PLIN2 dysregulates insulin secretion and [Ca]_j_ in INS1 cells. (a-c) (a) Representative confocal images of INS1 cells transfected with SiPLIN2 (SiPLIN2) and those treated with Scramble control (Scr) stained with Bodipy 493 (green) and DAPI (blue). Scale bar = 10 μm. (b) Violin plot for size of an individual lipid droplet (LD) expressed as area. Median and quartiles are indicated. (c) Cumulative frequency (%) of (b). Area of 69 LDs from 14 cells treated by Scr and 38 LDs from 13 cells treated by SiPLIN2 were measured. Representative data from 3 independent experiments. (d) Insulin secretion measured by static incubation at indicated concentrations of glucose for 1 h in INS1 cells transfected by Scr and SiPLIN2 with or without 24 h incubation in 0.2 mM palmitic acids (PA). Insulin secreted was corrected for mg protein and expressed taking the average of Scr treated at 2.5 mM as 1. Data combines three experiments each in triplicates. (e) Stimulation index determined as the ratio of average insulin secretion at 12 mM glucose over 2.5 mM glucose for each experiment. Each dot represents one experiment. (f) Total insulin contents for (d). Data were corrected for mg protein and expressed taking the average of Scr treated at 2.5 mM as 1. (g) Insulin secretion at 2.5 mM glucose as in (d) except for expressed as % total insulin contents. n=6. (h-i) Fura-2 Ca^2+^ transients in INS1 cells in response to 30 mM glucose was measured in 4 cover slips for Scr and 5 for SiPLIN2. (h) Glucose response calculated as Fura-2 340/380 as in methods. (i) Representative average tracings. n= 8 cells for Scr and 14 cells for SiPLIN2 per cover slide. Supplementary Fig. 3d shows tracing of an individual cell. All data are mean ± s.e.m. except for (b). *; p < 0.05 by Student’s t test.

Similar to islets from βKO on HFD, PLIN2 deficiency in INS1 cells reduced OCR in response to glucose, ATP production, and maximal respiration (Fig. 4a-d). Interestingly, the proton leak was elevated in SiPLIN2 treated INS1 cells that showed increase in basal insulin (Fig. 4e). In contrast, βKO islets did not show increase in basal insulin or increase in the proton leak (Fig. 2a, Supplementary Fig. 2a). Recently, the mitochondrial proton leak was proposed to trigger non-secretagogue insulin secretion under nutritional load through the activation of mitochondrial permeability transition pore (24). Thus, the increase in proton leak in SiPLIN2 treated INS1 cells may be a potential mechanism responsible for the increase in basal insulin secretion. As accelerated glucose metabolism is another mechanism proposed to increase insulin secretion at low glucose after nutritional stress (20, 22), we tested whether the inhibition of glucokinase by overnight incubation with D-Mannoheptulose reduces basal insulin secretion in SiPLIN2 treated INS1 cells. 0.5 mM D-Mannoheptulose appropriately reduced insulin secretion at 12 mM glucose in control, while basal insulin secretion in SiPLIN2 INS1 cells was decreased by 19% indicating that glucose metabolism only partly accounts for high basal insulin secretion in SiPLIN2 treated INS1 cells (Fig. 4f). Further, the high level of basal insulin secretion in SiPLIN2 INS1 cells did not depend on Ca^2+^ influx since nifedipine did not reduce insulin secretion at low glucose in SiPLIN2 INS1 cells (Fig. 4g). Mitochondrial complexes I, IV and V were reduced in SiPLIN2 treated INS1 cultured without chronic PA exposure (Fig. 4h). However, the exposure to 0.2 mM PA for 24 h did not affect the expression of mitochondrial complexes in Scr treated INS1 cells nor worsen reduction of complexes in siPLIN2 treated INS1 cells compared with PA untreated counterpart (Fig. 4h, Supplementary Fig. 3e). Mitochondrial DNA was significantly reduced in SiPLIN2 transfected INS1 cells indicating that mitochondrial mass is reduced after PLIN2 down-regulation (Fig. 4i). Collectively, down-regulation of PLIN2 in INS1 cells impaired mitochondrial function and integrity sharing the reduction of OCR and OXPHOS complex V with mouse βKO islet on HFD.

**Figure 4.**
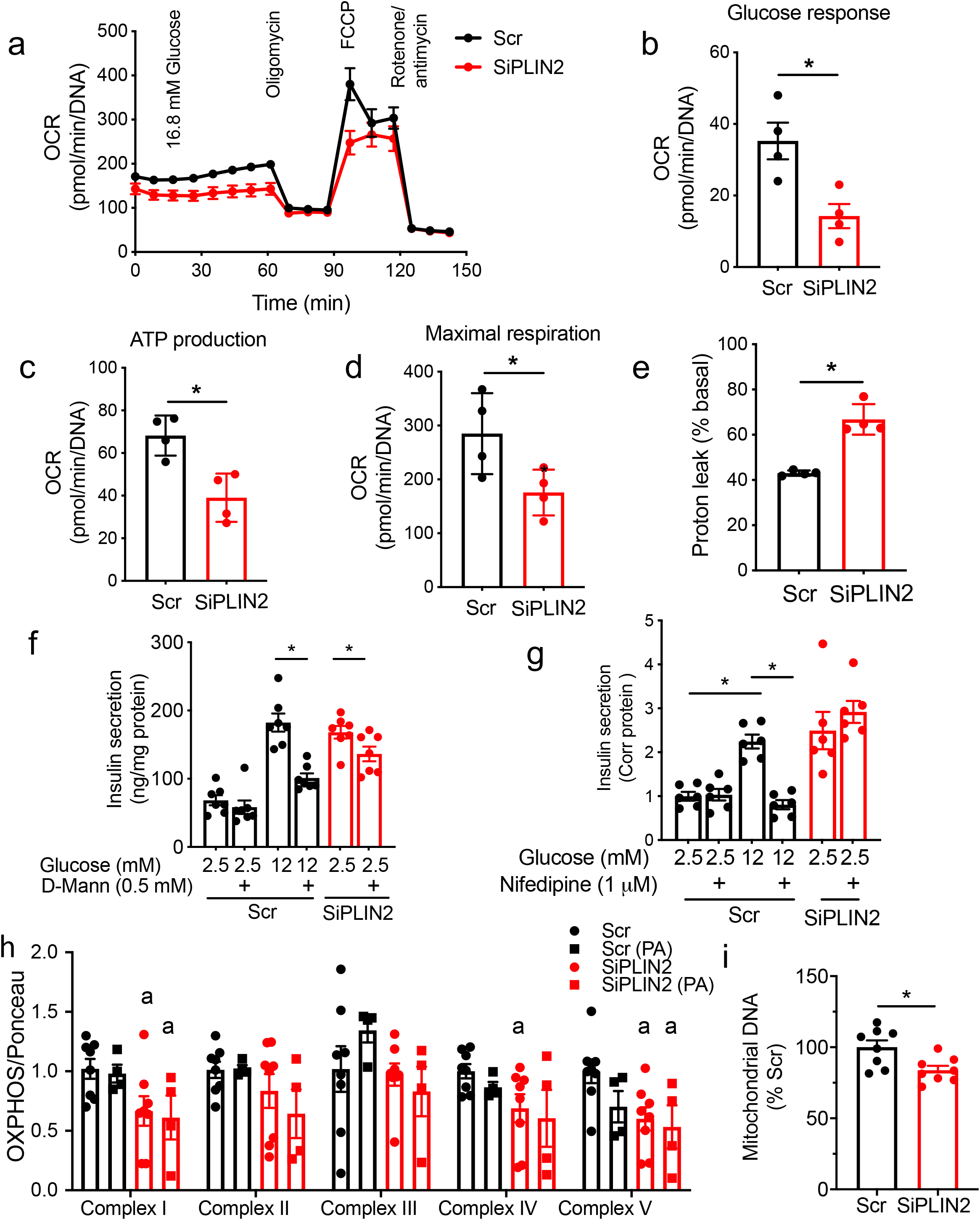
Down-regulation of PLIN2 causes mitochondrial dysfunction in INS1 cells. (a-e) Oxygen consumption rate (OCR) measured by Seahorse Metabolic analyzer and corrected for DNA contents in Scr and SiPLIN2 treated INS1 cells. (a) OCR profile representative of four experiments each performed in quadruplicates. (b) Glucose response, (c) ATP production, (d) maximal respiration, and (e) %proton leak. (b-e) Each dot is average value of one experiment. n=4. (f-g) Insulin secretion measured by static incubation and corrected for protein contents (f) with addition of 0.5 mM D-Mannoheptulose overnight and (g) 1 μM nifedipine for 2 h as in methods. n=6, representative of three experiments. *; *p*< 0.05 by Student’s t test. (h) Western blot of OXPHOS complex proteins were performed in protein lysate of INS1 cells transfected by Scr and SiPLIN2 treated with or without 0.2 mM palmitic acids (PA) for 24 h prior to harvest. Densitometry data were corrected for Ponceau S staining and expressed taking average value of Scr without PA as 1 for each complex. n=8 for no PA and 4 for PA treated INS1 cells. a: *p*< 0.05 vs Scr without PA. A representative blot in Supplementary Fig. 3e. (i) qPCR probed DNA from Scr and SiPLIN2 treated INS1 cells for mitochondrial DNA (mt-ND6) and nuclear DNA (β actin). Expression of mt-ND6 was corrected for β actin and the average of value for Scr was taken as 100%. n=8 combined from three independent experiments. All data are mean ± s.e.m.. *; *p*< 0.05.

### PLIN2 downregulation in INS1 cells increases trafficking of FA to mitochondria

To address why down-regulation of PLIN2 affects mitochondria, we investigated how PLIN2 deficiency alters FA distribution within cells by incubating SiPLIN2 treated INS1 cells overnight with Bodipy 558/568 C12 (Bodipy C12), a fluorescent oleic acid (OA) analog that is preferentially incorporated into TG (Fig. 5a) (25, 26). In Scr treated cells, Bodipy C12 was highly incorporated in LD defined by Bodipy 493 (Fig. 5a, b). In contrast, SiPLIN2 caused a significant reduction in the intensity of Bodipy C12 within each LD and % of Bodipy C12 distributed to LDs in each cell (Fig. 5a-c) in agreement with a previous study in MIN6 cells that demonstrated that PLIN2 down-regulation reduces [^3^H]OA incorporation into TG measured by thin layer chromatography (12). Since OA uptake measured using [^3^H]OA did not differ between Scr and SiPLIN2 treated cells (Fig. 5d), there likely is the more FA distributed outside of LDs in SiPLIN2 cells. Indeed, the high proportion of Bodipy C12 co-localized with a mitochondrial marker, heat shock protein 60 (HSP60) indicating that a larger proportion of FA is targeted to mitochondria in PLIN2 deficient INS1 cells (Fig. 5e). We also noted the accelerated fragmentation of mitochondria when SiPLIN2 treated INS1 cells were treated with 0.4 mM OA, that was reported to promote TG formation with minimum cytotoxicity in INS1 cells (Fig. 5f-h) (19). An aspect ratio and a form factor were both decreased in SiPLIN2 treated INS1 cells indicating mitochondria are rounder and fragmented, changes also seen in INS1 cells under nutritional stress (Fig. 5h, supplementary Fig. 3f) (27, 28).

**Figure 5.**
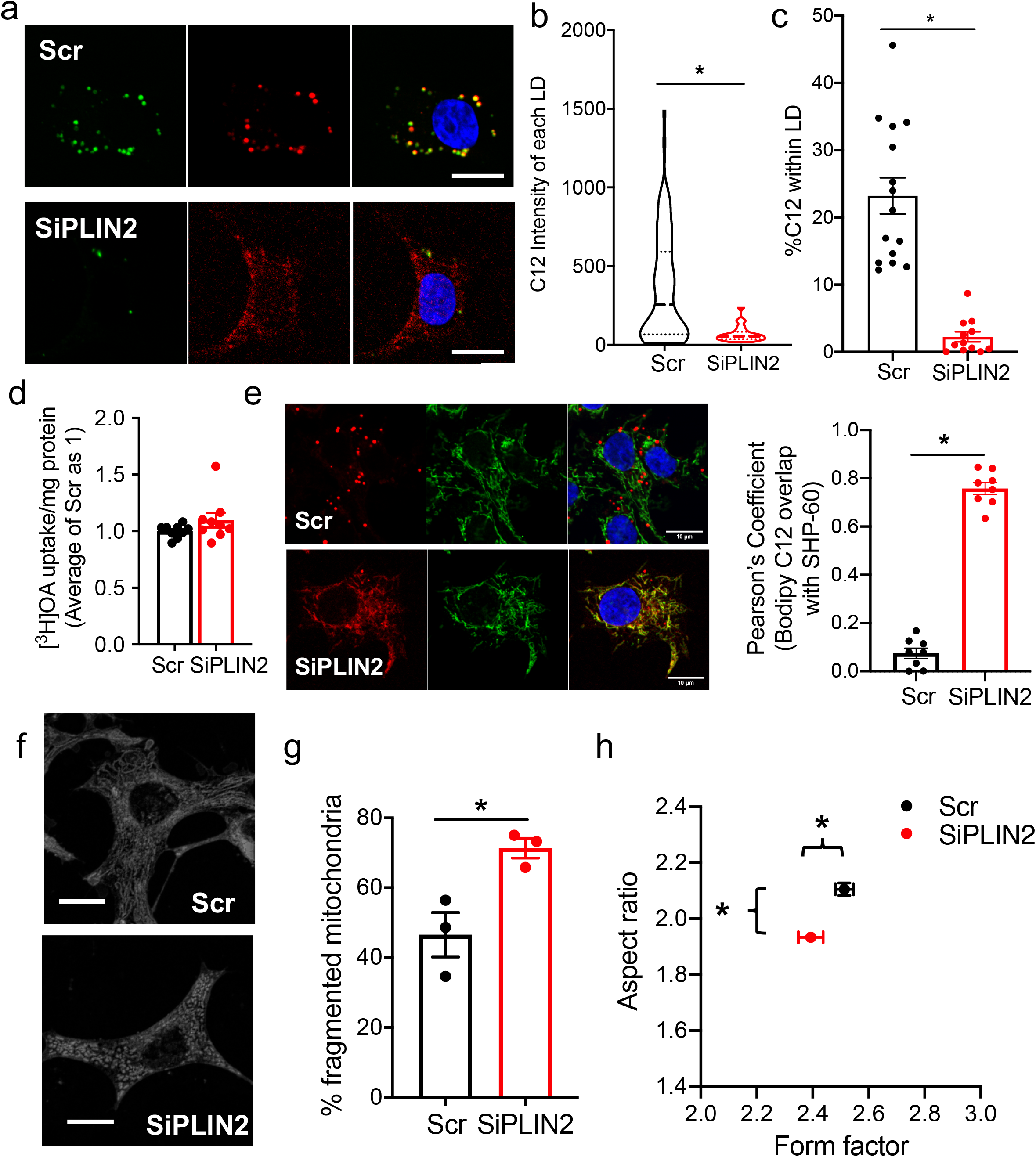
PLIN2 down-regulation alters distribution of Bodipy C12 in INS1 cells. (a) Representative confocal images of INS1 cells transfected with SiPLIN2 (SiPLIN2) and scramble control (Scr) and metabolically labeled with Bodipy C12 (red, middle panel) overnight followed by staining with Bodipy 493 (green, left panel) and DAPI (blue). Merged pictures on the right. (b) Violin plot of integrated intensity of Bodipy C12 signal within each individual lipid droplet (LD) defined by Bodipy 493. Median and quartiles are indicated. 344 LDs from 15 cells transfected with Scr and 149 LDs from 16 cells transfected with SiPLIN2 were measured. (c) Proportion of Bodipy C12 signal found in LDs defined by Bodipy 493 in 15 cells transfected with Scr and 12 cells transfected with SiPLIN2 was measured. (b, c) Representative data from 3 independent experiments. (d) [^3^H]OA uptake after overnight incubation of Scr and SiPLIN2 treated INS1 cells was corrected for protein contents and expressed taking the average value of Scr as 1. n=9. (e) Representative confocal images of Scr and SiPLIN2 treated INS1 cells metabolically labeled with Bodipy C12 (red, left panel) overnight followed by immunostaining by HSP60 (green, middle panel). Merged pictures on the right. Pearson’s coefficient for Bodipy C12 and HSP60 was calculated as in methods. n= 8. Representative data from 3 independent experiments. (f) Representative MitoTracker Deep Red images of Scr and SiPLIN2 INS1 cells. (g) % of fragmented mitochondrial defined as aspect ratio below 2 in three independent experiments. Number of mitochondria counted for Scr was exp1=74, exp2=55, exp3= 81, and for SiPLIN2 was exp1=73, exp2=41, exp3= 40. (h) An aspect ratio and a form factor for all mitochondria counted in three independent experiments. n= 210 for Scr and 154 for siPLIN2. See Supplementary Fig. 3f for violin plot data. All scale bars are 10 μm. Data represent mean ± s.e.m. for all except (b). *; *p*<0.05 by student’s t test.

### PLIN2 down-regulation in INS1 cells impacts nutrient metabolism

To address whether the increased targeting of FA to mitochondria alters FA metabolism in SiPLIN2 treated INS1 cells, FA oxidation (FAO) was assessed by measuring the production of [^3^H]water from [^3^H]OA and found to be reduced in SiPLIN2 INS1 cells (Fig. 6a). As the increase in acylcarnitines due to incomplete FAO is proposed to contribute to β cell dysfunction (29–31), we analyzed 81 metabolites that are important for energy homeostasis including 13 acylcarnitines in SiPLIN2 treated and control INS1 cells under regular culture condition (Supplementary Table 1). 41 metabolites including 8 carnitines showed statistically significant differences between SiPLIN2 and control INS1 cells with a clear separation of two groups by principal component analysis (PCA, Supplementary Fig. 4, Supplementary Table 2). For acylcarnitines with an even number of acyl carbons, there was overrepresentation of mid chain C12:0 acylcarnitine with reduction of both short chain (C2:0 and C4:0) and long chain (C16:0 and C18:0) acylcarnitines consistent with reduced oxidation of mid chain FA in SiPLIN2 treated INS1 cells (Fig. 6b). For odd chain acylcarnitines, C5:0 was increased but C3:0 was reduced in SiPLIN2 treated INS1 cells (Fig. 6c). In addition, the number of glucose metabolites in glycolysis, TCA cycle, and pentose phosphate pathway (PPP) were reduced in SiPLIN2 treated INS1 cells (Fig. 6d), which could also contribute to impaired GSIS (32). Amino acids primarily metabolized to pyruvate were reduced but those primarily metabolized to acetyl-CoA and intermediates of TCA cycle were increased in SiPLIN2 treated INS1 cells (Fig. 6e). This likely reflect reduced utilization of amino acids by the TCA cycle considering that these amino acids were supplied in culture medium. Although the current result is only a snapshot of nutrient metabolism, it is of note that a wide range of metabolites connected with glucose metabolism and important for insulin secretion were altered in PLIN2 deficient INS1 cells. Other significant changes in SiPLIN2 treated INS1 cells included an increase in glutathione and carnosine that are known to reduce oxidative stress (Fig. 6f) (33, 34). This may compensate for mitochondrial dysfunction and explain the lack of increase in 4-Hydroxynonenal, a marker of oxidative stress, in SiPLIN2 treated INS1 cells (data not shown).

**Figure 6.**
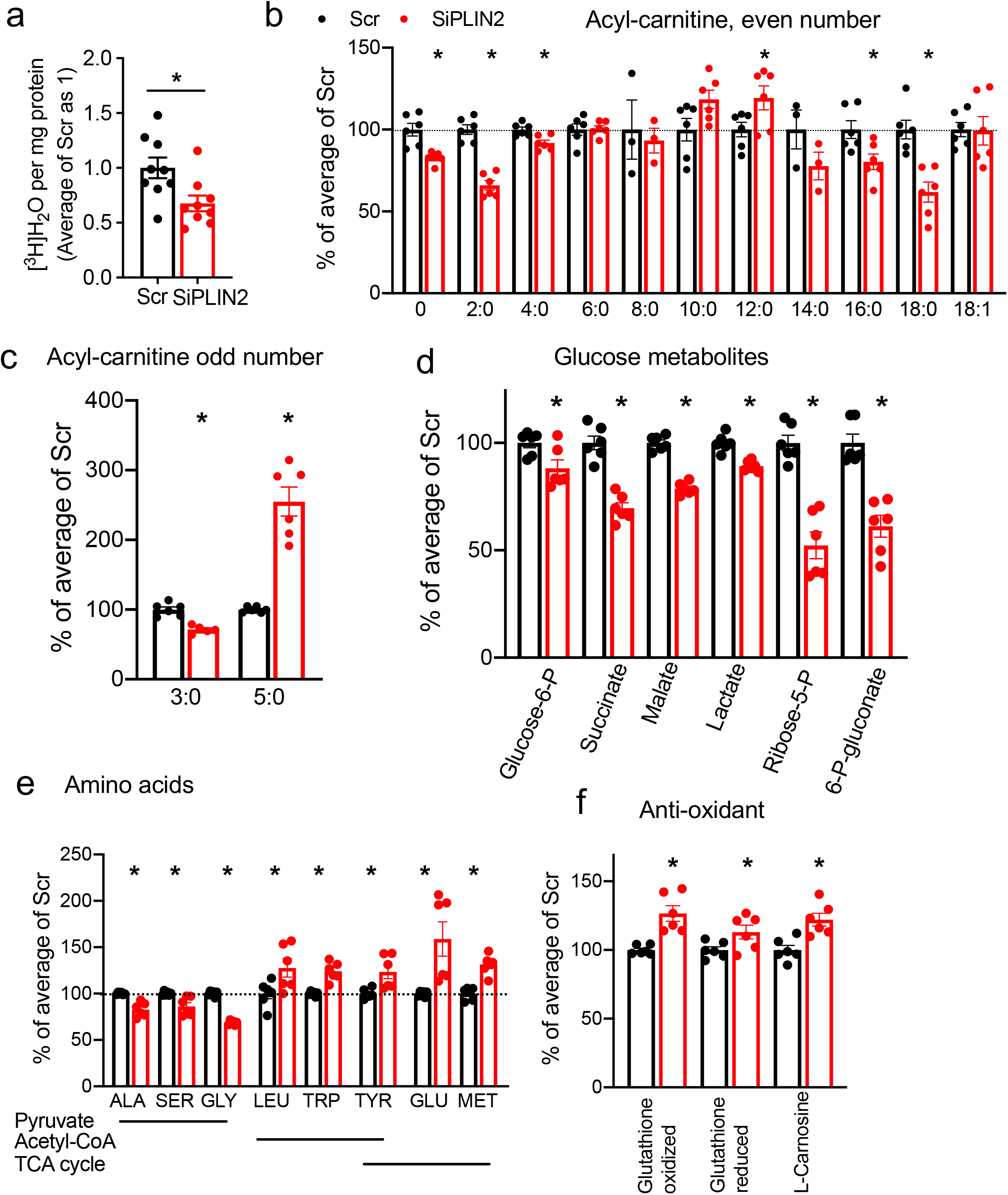
PLIN2 down-regulation alters nutrient metabolism in INS1 cells. (a) [^3^H]water generated after an overnight incubation with [^3^H]OA in INS1 cells transfected with scramble control (Scr) and SiRNA targeting Plin2 (SiPlin2) corrected for protein contents and expressed taking the average value of Scr as 1. n=9. (b-f) Relative abundance of metabolites measured by LC-MS/MS in Scr (•) and SiPLIN2 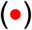 treated INS1 cells expressed taking the average value of Scr as 100%. n=6 except for C14:0 in (b) that is n=3. Data are mean ± s.e.m.. *; p < 0.05 by student’s t test.

### PLIN2 down-regulation does not reduce ER stress in β cell models

Previously, it was reported that whole body PLIN2 KO mice with an Akita insulin mutation preserve β cell mass compared with Akita mice with intact PLIN2 (13). Thus, we tested whether PLIN2 deficiency in our β cell models reduced ER stress provoked by tunicamycin. While tunicamycin significantly increased *Hspa5* and *Ddit3* expression in control INS1 cells, down-regulation of *Plin2* by siRNA did not reduce the expression of *Hspa5* and *Ddit3* significantly in the presence and absence of tunicamycin (Fig. 7a-c). Islets from βKO mice on regular chow were not protected against the tunicamycin induced rise in *Hspa5* and *Ddit3* expression (Fig. 7d-f). Tunicamycin reduced *Plin2* at mRNA levels in both INS1 cells and mouse islets (Fig. 7c, f), which was somewhat the opposite of a reported increase in *Plin2* level in MIN6 cells and mouse islets after tunicamycin exposure (13). Thus, we assessed how tunicamycin alters PLIN2 expression in human islets. When human islets from non-diabetic donors were incubated with tunicamycin or OA overnight, tunicamycin clearly provoked ER stress as evidenced by the increase of *HSPA5, DDIT3,* and *XBP1s* (Fig. 8a-c). OA also increased *DDIT3* and *XBP1s,* but not *HSPA5* (Fig. 8a-c). However, the increase of PLIN2 both at the mRNA and protein levels was apparent only in OA treated human islets but not with tunicamycin treatment indicating that ER stress is not a major driver to increase PLIN2 in human islets (Fig. 8d, e).

**Figure 7.**
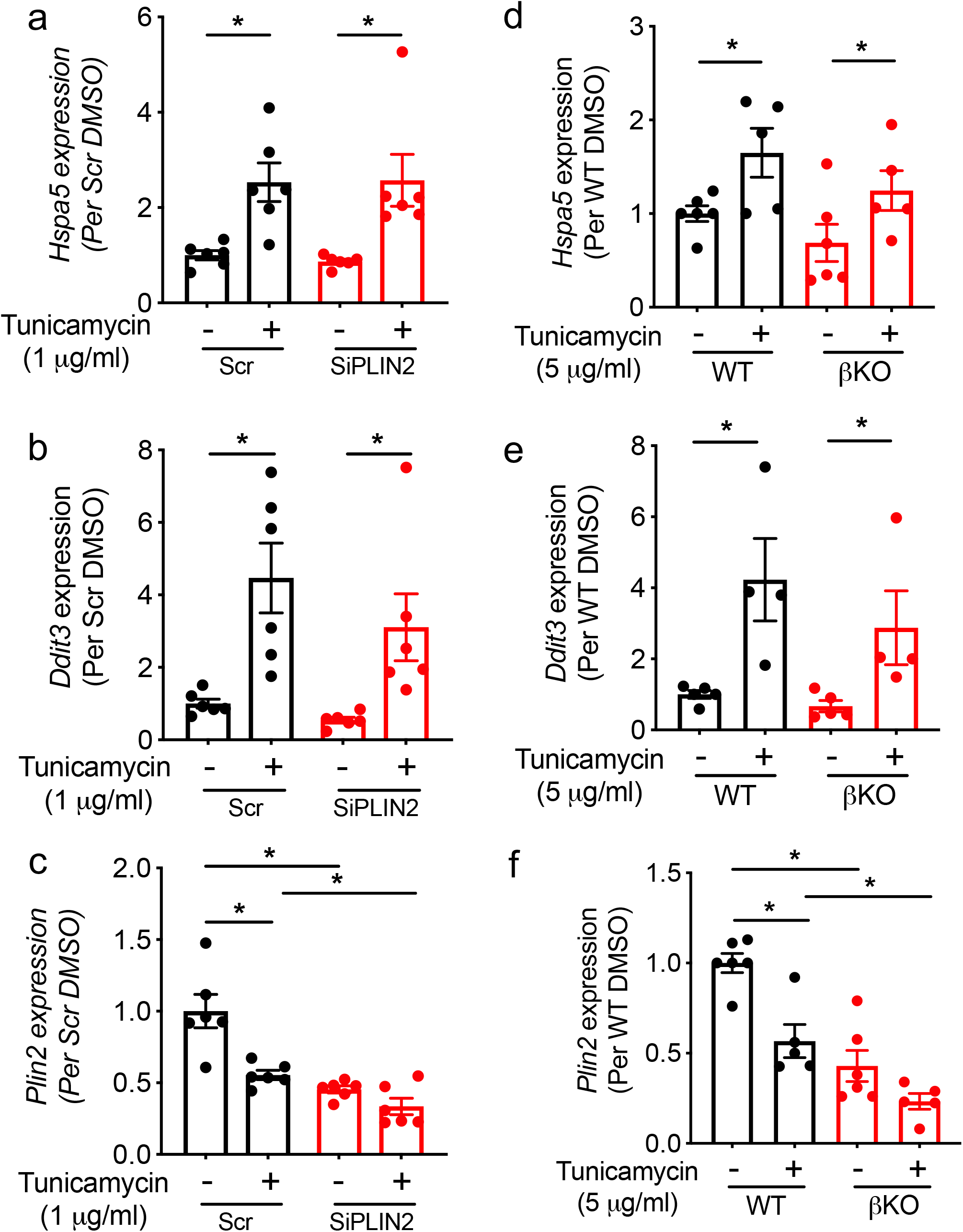
PLIN2 down-regulation does not reduce ER stress in INS1 cells and mouse islets. (a-c) INS1 cells transfected with scramble control (Scr) and SiPLIN2 were treated with DMSO or tunicamycin for 6 h prior to harvest and the expression of *Hspa5* (a), *Ddit3* (b), and *Plin2* (c) were measured by qPCR using *Ppib* as internal control. n=6. (d-f) Expression of *Hspa5* (d), *Ddit3(e),* and *Plin2* (f) in islets of β cell specific PLIN2 KO (βKO) and WT littermate mice on regular chow cultured with DMSO or tunicamycin for 6 h prior to harvest. *Ppib* as internal control. n= 4-6 for WT_fl_ and βKO. Data are expressed taking the average value of Scr or WT as 1 and mean ± s.e.m.. *; *p*< 0.05 by student’s t test.

**Figure 8.**
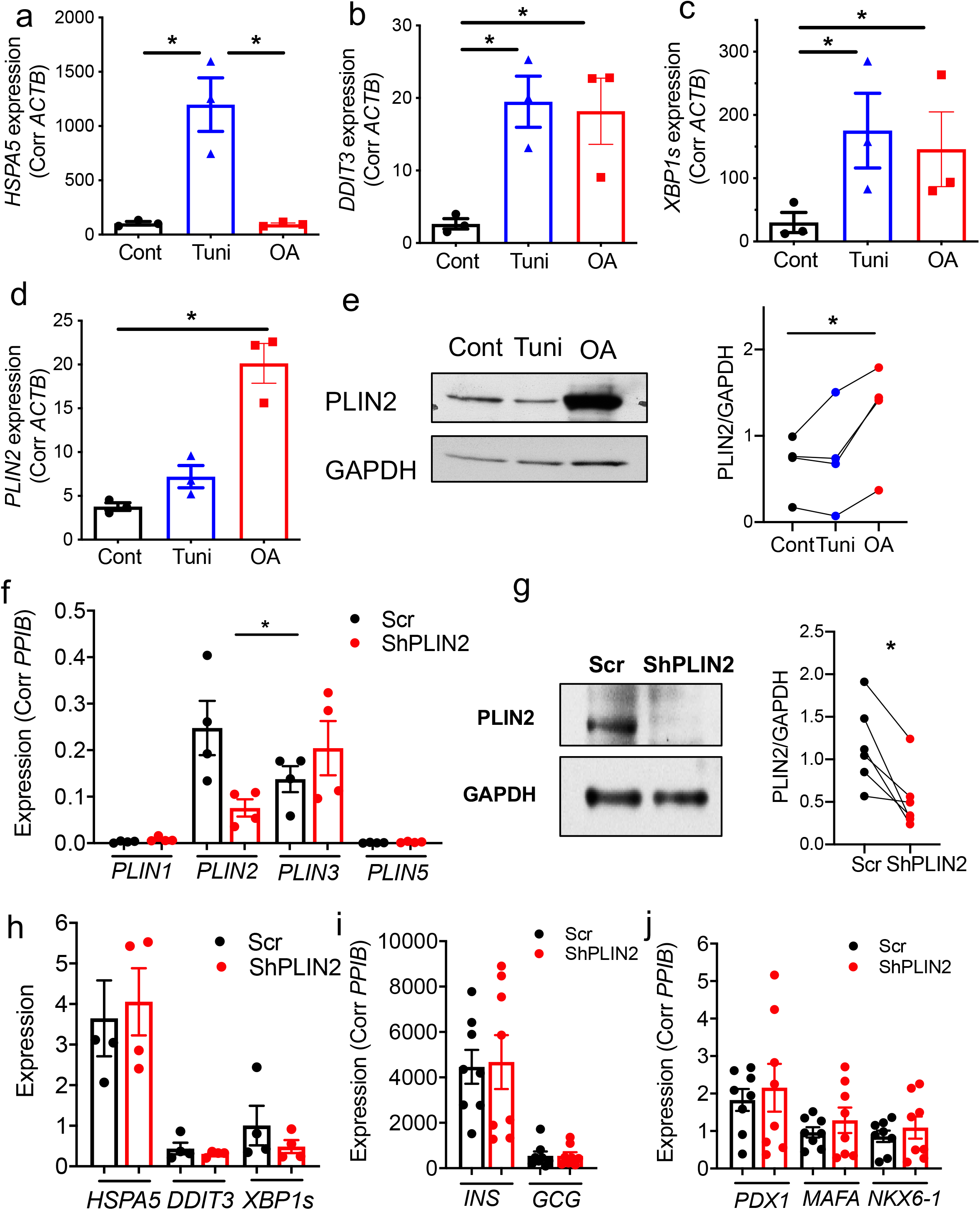
ER stress is not the major driver of PLIN2 expression in human islets. (a-d) qPCR compared the expression of *HSPA5* (a), *DDIT3* (b), *XBP1S* (c), and *PLIN2* (d) in human islets from non-diabetic donors incubated with 10 μg/ml tunicamycin (tuni) or 0.4 m OA overnight. Data are expressed using *ACTB* as an internal control. n=3 donors. (e) A representative blot and densitometry of Western blot comparing the expression of PLIN2 in nondiabetic human islets treated as for (a-d). Data are corrected for GAPDH. Each dot represents a single donor and data from the same donor is connected by a line. n=4 donors. (f) qPCR assessed the expression of PLINs in human pseudoislets treated with lenti-shPLIN2 (ShPLIN2) and lenti-ShScr (Scr). Expression was corrected using *PPIB* as internal control. (h) A representative blot and densitometry of Western blot comparing the expression of PLIN2 in ShPLIN2 and Scr human pseudoislets cultured with 0.5 mM OA 16 h prior to harvest. Data are corrected for GAPDH. n=6 donors. (h-j) qPCR assessed the expression ER stress markers (h), hormones (i), and markers of β cell maturation in human pseudoislets treated with Scr or shPLIN2. Expression was corrected by *PPIB* as an internal control for all except for *XBP1s* for which *HPRT1* was used. n=4 for h and n= 8 for i and j. All data are mean ± s.e.m. and each dot represents a single donor. *; *p*< 0.05 by student’s t test.

### PLIN2 deficiency blunts insulin secretion and reduces OXPHOS complex in human pseudoislets

Next, we down-regulated PLIN2 without changing the expression of other PLINs in human pseudoislets using lenti-ShRNA (Fig. 8f) (35). The reduction of PLIN2 was also confirmed by Western blot in shPLIN2 treated human pseudoislets (Fig. 8g). Down-regulation of PLIN2 did not change the expression of ER stress markers, *INS, GCG,* or maturation markers of β cells including *PDX1, MAFA,* and *NKX6-1* in human pseudoislets (Fig. 8h-j). However, both glucose-stimulated and KCL-stimulated insulin secretion was significantly reduced to 20% and 30% of control in shPLIN2 human pseudoislets cultured in 10% FBS CMRL medium (Fig. 9a-c). There also was moderate reduction of basal insulin secretion to 49% of Scr control in shPLIN2 human pseudoislets (Fig. 9d). As there was mild but significant increase in total insulin contents in shPLIN2 pseudoislets (1.76 ± 0.20 fold of Scr control, mean ± s.e.m, *p*<0.05, n=9), SI was also compared as a parameter independent from insulin contents. Notably, the SIs for first phase (Fig. 9e), second phase (Fig. 9f), and the entire GSIS (Fig. 9g) were all reduced in shPLIN2 human pseudoislets. As parts of the mitochondrial complexes were reduced in βKO islets from mice on HFD and SiPLIN2 treated INS1 cells, mitochondrial complexes were compared and was found to be reduced for Complex III in shPLIN2 treated human pseudoislets (Fig. 9h).

**Figure 9.**
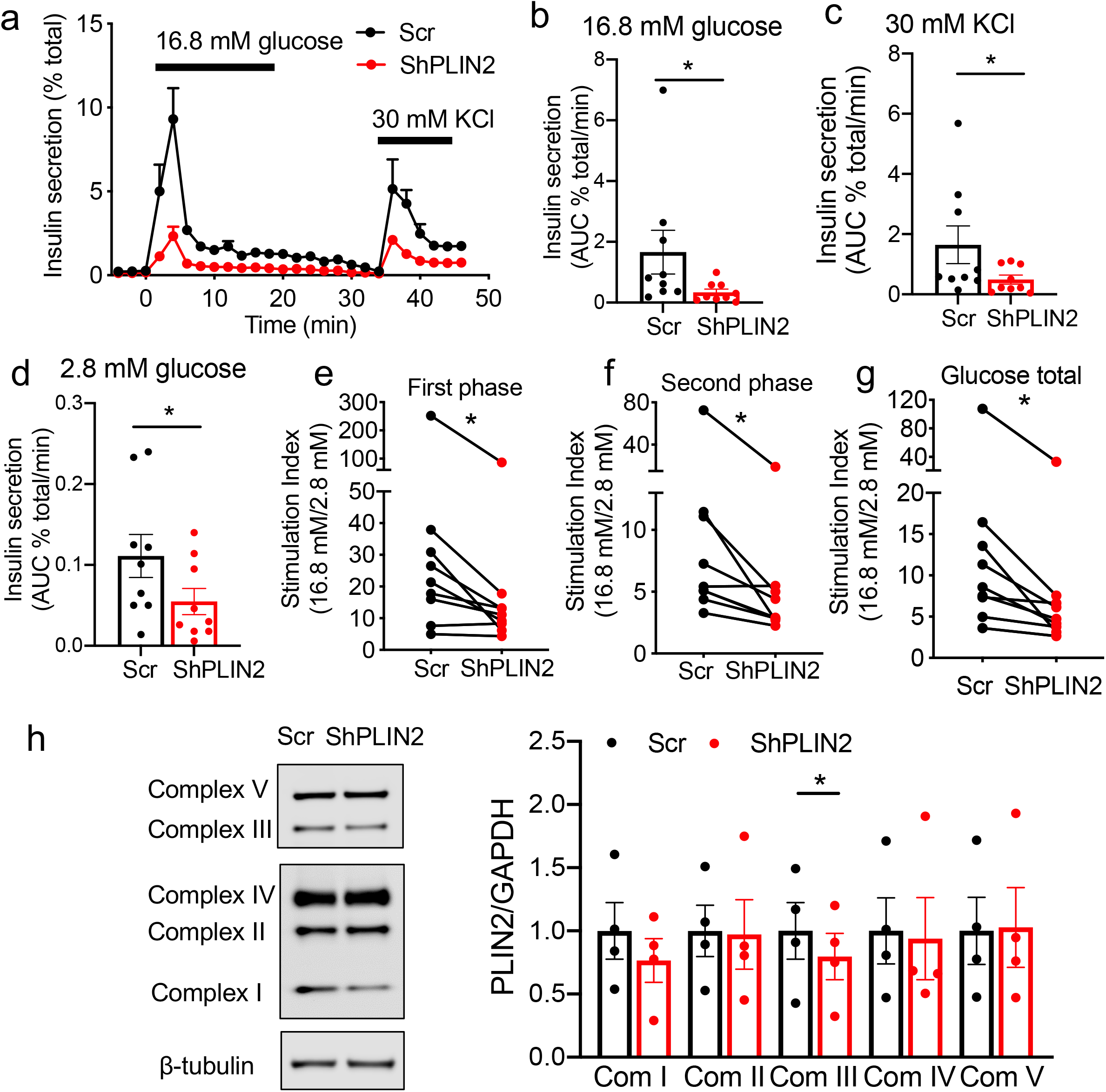
Insulin secretion of human pseudoislets transduced by ShPLIN2. (a) A representative profile of insulin secretion by perifusion of human pseudoislets transduced by lenti-shPLIN2 (ShPLIN2) and lenti-ShScr (Scr) in response to 16.7 mM glucose and 30 mM KCl. Data are expressed as % of total insulin. Mean ± s.e.m of duplicates from a single donor. (b-d) Area under the curve (AUC) representing the average insulin secreted over each indicated treatment period. (b) 16.8 mM glucose ramp, (c) 30 mM KCl, and (d) 2.8 mM glucose. (e-g) Stimulation index during first phase (e), second phase (f) and the entire 16.8 mM glucose ramp (g) were obtained as in methods. Each dot represents a single donor and a line connects data from the same donor. n=9 donors. (h) Western blot of OXPHOS complexes in protein lysate of Scr (•) and ShPLIN2 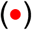 treated human pseudoislets cultured with 0.5 mM oleic acid for 16 h prior to harvest. Due to difference in intensity of bands, light exposure is shown for complex V and III and dark exposure is shown for the rest of complexes in the representative blot. Densitometry data were corrected for β-tubulin and expressed taking average value of Scr as 1 for each complex. n=4 donors. All data are mean ± s.e.m.. *; *p*<0.05 by Student’s t test.

## Discussion

Down-regulation of PLIN2 negatively affected GSIS in βKO mice fed HFD, in SiPLIN2 treated INS1 cells, and in ShPLIN2 treated human pseudoislets indicating that PLIN2 is indispensable for normal GSIS and that the reduction of PLIN2 accelerates β cell dysfunction under nutritional stress. Blunting of GSIS was associated with reduced OCR in response to glucose in islets from HFD fed βKO mice and in SiPLIN2 treated INS1 cells pointing to a defect up-stream of oxidative phosphorylation. Mitochondrial complex proteins were reduced in βKO islets, in SiPLIN2 treated INS1 cells, and in shPLIN2 treated human pseudoislets indicating that the integrity of mitochondria is impaired when PLIN2 is down-regulated in β cells. The increase in proton leak and fragmentation of mitochondria with OA loading in SiPLIN2 treated INS1 cells further indicates that PLIN2 coated LDs are protective for mitochondria in β cells. PLIN2 deficiency in INS1 cells resulted in reduced levels of FA incorporated into LD and increased trafficking of FA to mitochondria, which might have negatively affected mitochondrial function.

In response to nutritional lipid loading, PLIN2 increased in MIN6 cells, rat islets, mouse islets, and human islets indicating that PLIN2 is the key PLIN isoform to support expansion of the LD pool during nutrient overload in pancreatic islets (8, 9, 12). PLIN2 has been proposed to increase LD lipid levels by reducing but not completely blocking lipolysis mediated by adipose triglyceride lipase (ATGL) (11, 36). Lipolysis corrected for TG accumulation was reduced in HEK293 and MIN6 cells overexpressing PLIN2 (9, 37). Thus, PLIN2 allows expansion of the LD pool in β cells while maintaining a certain level of lipolysis, which is important for β cells that utilize lipolytic metabolites for insulin secretion (4, 17, 38). In agreement, overexpression of PLIN2 in MIN6 cells increased the TG pool with little effect on GSIS, thereby showing the ability of PLIN2 to expand the inert neutral lipid pool in β cells (12). PLIN2 is stabilized when it interacts with LDs; consequently, PLIN2 enables β cells to quickly adapt to an increase in neutral lipid synthesis by reducing lipolysis (39, 40). The impairment of GSIS in βKO mice on HFD and in SiPLIN2 INS1 cells exposed to PA indicates that up-regulation of PLIN2 preserves β cell function when nutritional stress drives expansion of the LD pool. In INS1 cells and in human pseudoislets that possess prominent LDs, impaired insulin secretion was seen even under regular culture conditions after PLIN2 down-regulation further supporting a protective role of LDs for β cell function (17).

Mitochondria can be spatially and functionally associated with LDs through LD formation and FAO (25, 41, 42). Nguyen *et al.* showed that LD formation buffers FA released by autophagosomal digestion of phospholipids in nutrient deprived MEF. In their study, LDs prevented mitochondrial dysfunction by limiting exposure of mitochondria to acylcarnitine generated from FA (43). In SiPLIN2 treated INS1 cells, reduction of LD formation increased the distribution of FA to mitochondria, which might have triggered the impairment in complete FAO leading to an increase in mid-chain acylcarnitines. In β cells, a significant proportion of glucose and FA are directed to TG synthesis within minutes of exposure to glucose indicating that β cells actively synthesize neutral lipids concurrent with insulin secretion (44, 45). Thus, the formation of PLIN2 coated LDs likely has an important function in protecting mitochondria in β cells, a protection that could become even more important when β cells are exposed to excess FA, as seen in βKO mice fed HFD and in SiPLIN2 treated INS1 cells cultured with PA.

Acylcarnitines, intermediate metabolites of FAO, are known to be increased when FA influx to mitochondria surpasses energy demands and the capacity of the TCA cycle (46). Both long and medium chain acylcarnitines were shown to impair mitochondrial function and reduce insulin secretion in mouse islets (30, 31). Recent evidence indicates that acylcarnitines are increased in β cells in T2D and may contribute to β cell dysfunction. Aichler *et al* reported the accumulation of stearoyl-carnitine (C18:0) in islets affected by T2D and in mouse models of T2D (30). In our study of SiPLIN2 treated INS1 cells under regular culture condition, a mid chain acylcarnitine (C12:0) was increased while both long chain acylcarnitines (C16:0, C18:0) and short chain acylcarnitines (C2:0, C4:0) were reduced. Although the reduction of long chain acylcarnitines may appear contradictory to the increase in long chain acylcarnitines reported in T2D islets (30), the discrepancy could be due to the limited availability of carnitine in culture systems. Using mouse islets in culture, Doliba *et al* showed that high glucose and FA markedly increase 3-hydroxytetradecaenoyl-carnitine (3-OH-C14), which is known to inhibit oxidative phosphorylation in rat heart mitochondria (29). In Doliba’s study, the increase in 3-OH-C14 was also associated with the reduction of C16:0 and C2:0 carnitines in mouse islets cultured with high glucose and PA (29). Alternatively, peroxisomal FAO might have contributed to the reduction of long chain acylcarnitine in SiPLIN2 treated INS1 cells (47). Also, the study of circulatory acylcarnitines indicated that the increase in mid chain acylcarnitines may precede the increase in long chain acylcarnitines during the development of beta cell dysfunction (31). Thus, the accumulation of mid chain acylcarnitine might have mediated mitochondrial dysfunction in PLIN2 deficient β cells.

While the necessity of PLIN2 for GSIS was demonstrated in all three models (12), the rise of basal insulin was seen only in SiPLIN2 treated INS1 cells. The increase in basal insulin is considered to be an early adaptation to nutritional stress and is seen in human islets, mouse islets, and β cell lines exposed to elevated glucose and FA (20–23). Compared with mouse islets and human islets, INS1 cells may be more susceptible to stress from the reduction of PLIN2 as INS1 cells have very low expression of PLIN3, another PLIN with functional redundancy with PLIN2 (18). Alternatively, the short period (72 h) of intervention to reduce PLIN2 in INS1 cells may account for difference in basal insulin secretion compared with mouse islets (6 months) and human pseudoislets (7 days) that were subjected to longer periods of PLIN2 reduction. It is also important to note that β cell dysfunction in βKO mouse on HFD appeared to be mild and insufficient to impair glucose tolerance. Unlike human T2D that is known to be progressive, hyperglycemia in C57BL/6J mice on HFD remains modest as they adapt by increasing β cell mass (16). Morphometry showed that β cell area was similar between βKO and WT littermate mice implying that PLIN2 deficiency does not affect the adaptive increase in β cell mass. Also, mouse β cells do not make large LDs as seen in human β cells and INS1 cells (17), which may make the impact of losing PLIN2 less severe. Nevertheless, our data clearly indicates that the loss of PLIN2 in β cells does not confer any protection against development of the modest hyperglycemia observed in mice on HFD.

Worsening of β cell function in PLIN2 deficient β cells under HF conditions and PA exposure is in contrast to previous studies of liver and adipose tissue that showed PLIN2 down-regulation reduced adversity associated with nutritional stress. ASO mediated down-regulation of PLIN2 improved insulin sensitivity in C57BL/6J mice fed HFD and in *ob/ob* mice (48). McManaman *et al.* found that Δ5PLIN2 whole body KO mice fed HFD were protected against obesity with increased beiging of adipocytes, reduced hepatosteatosis, lower adipose inflammation, and increased mobility (49). One proposed mechanism for the improvement in hepatosteatosis in PLIN2 KO mice is reduced activation of SREBP-1 and −2 in the ER decreasing TG and cholesterol synthesis. The increase in beiging in Δ5PLIN2 whole body KO was prominent in mice fed high sucrose and correlated with hepatic FGF21 expression, implicating that the loss of hepatic PLIN2 stimulates FGF21 production (50). Given that both hepatocytes and adipocytes have a specialized function in TG metabolism with large LDs and high expression of multiple PLINs, the different consequence of PLIN2 depletion is not necessarily surprising and highlights a cell type specific role of PLIN2.

It was previously reported that ER stress drives PLIN2 expression in mouse islets and that the reduction of PLIN2 protects β cells from ER stress by increasing autophagy (13). Akita mice with Δ2-3PLIN2 whole body KO were less hyperglycemic than Akita mice with intact PLIN2 (13). However, the reduction of PLIN2 was neutral to tunicamycin induced ER stress and caused a mild increase in *Ddit3* in islets of high fat fed βKO mice in the current study. It is plausible that the loss of PLIN2 in β cells confers protection under severe ER stress and extreme hyperglycemia as seen in Akita mice. It is also possible that Δ2-3PLIN2 whole body KO mice may be less hyperglycemic due to an extra islet effect of PLIN2 deletion, considering that carbohydrates strongly drive FGF21 expression in PLIN2 deficient liver (50). Ultimately, it will be important to determine whether the loss of PLIN2 confers protection to ER homeostasis in human β cells during the development of T2D. Although down-regulation of PLIN2 in human pseudoislets was not sufficient to alter markers of ER stress under regular culture condition, it will be important to address whether PLIN2 and LD formation reduce ER stress induced by over nutrition in human islets.

The current study has several limitations. Our data collectively indicate that mitochondria are negatively affected by down-regulation of PLIN2 in β cells. However, further study should address whether mitochondrial dysfunction is solely responsible for the impairment in insulin secretion, especially in human islets after PLIN2 down-regulation. In human pseudoislets, the impairment of the dynamic secretion of insulin after PLIN2 down-regulation was not associated with an increase in markers of ER stress or a reduction in β cells markers. On the other hand, mitochondrial complex III was mildly, but significantly, reduced in ShPLIN2 human pseudoislets, indicating that impaired integrity of mitochondria contributes to impaired insulin secretion. However, we are yet to demonstrate the extent of impact of PLIN2 deficiency on multiple aspects of mitochondria in human β cells and to answer whether PLIN2 deficiency in human β cells increases FA distribution to mitochondria as seen in INS1 cells. It also should be noted that lenti-shPLIN2 we utilized is not specific for β cells and can reduce expression of PLIN2 in other human islet cells as well. As down-regulation of PLIN2 was reported not to affect viability of INS1 cells under FA exposure (51), we focused our study of PLIN2 down-regulation on β cell function. However, it remains to be determined whether PLIN2 deficiency in human beta cells accelerates beta cell demise under prolonged nutritional stress *in vivo* using a model such as transplantation of PLIN2 deficient human pseudoislets to immunodeficient mice on HFD, a model known to increase LD formation in human beta cells (10).

In conclusion, PLIN2 is indispensable for normal insulin secretion in HFD fed mice, in INS1 cells, and in human pseudoislets. A reduction in OXPHOS proteins was noted in all three models indicating that PLIN2 is important to maintain the integrity of mitochondria in β cells. The importance of PLIN2 for mitochondrial health was further demonstrated when the down-regulation of PLIN2 in INS1 cells increased FA distribution to mitochondria and caused multiple changes including reduced OCR, incomplete FAO, alterations in acylcarnitine, reduced glucose metabolism, and susceptibility of mitochondria to fragmentation. Thus, the PLIN2 coated LD is an important organelle to mitigate nutritional stress in β cells.

## Methods

### Animal studies

Experiments were performed in accordance with the Institutional Animal Care and Use Committee guidelines of University of Iowa. Mice with loxP flanking exon 5 of *Plin2* were previously described (49) and crossed with Ins1Cre mice (026801, Jackson Laboratory) on a C57BL/6J background (52). Mice were housed 5/cage in 12 h light-dark cycle at 22 °C, allowed free access to water, and fed regular rodent chow (7319 Teklad global diet) or HFD (45 kcal% fat, D12451, Research Diets). Mouse islets were isolated by collagenase P (Roche) digestion of the pancreas followed by Ficoll (GE healthcare life science) density centrifugation and cultured overnight in 11 mM glucose RPMI1640 supplemented with 10% FBS as described (53).

### GTT and ITT

GTT was done after 6 h fasting by loading 1 mg/g BW glucose i.p. and tail blood glucose was measured at time 0, 15, 30, 60, 90 and 120 min using a handheld glucometer. We also performed a study using 1.5 mg/g BW glucose i.p. and obtaining blood at time 0 and 15 min to measure serum insulin using ELISA (Alpco). ITT was performed after overnight fasting by 0.75 mU/g BW regular insulin i.p. and glucose was measured at time 0, 15, 30, 60, and 120 min.

### INS1 cells

823/13 cells (INS1 cells, a kind gift from Dr. Christopher Newgard (Duke University)) were maintained in RPMI1640 supplemented with 10 mM HEPES, 10%FBS, 2 mM L-glutamine, 1 mM sodium pyruvate, 50 μM β-mercaptoethanol, and penicillin+ streptomycin (INS1 medium). For down-regulation of PLIN2, cells were transfected with 30 nM of siRNA targeting PLIN2 (rn.Ri.Plin2.13.1 from Integrated DNA Technologies) using DharmaFECT1 transfection reagent as published (17). Non-targeting DsiRNA (Integrated DNA Technologies) was used as negative control. Experiments were performed 72 h after transfection.

### Static incubation

After 1 h pre-incubation in glucose free Krebs-Ringer buffer (KRB), INS1 cells or mouse islets (10 islets/1.5 ml tube) were incubated for 1 h in KRB containing the indicated concentration of glucose. Insulin secreted and insulin contents were measured with an ultrasensitive mouse Insulin ELISA and STELLUX Chemiluminescent rodent Insulin ELISA respectively (ALPCO). D-Mannoheptulose dissolved in DMSO was added overnight and continued through GSIS. Nifedipine dissolve in DMSO was added 1 h prior to GSIS.

### mRNA and quantitative PCR

RNA was isolated from INS1 cells using RNeasy kit (Qiagen) and from islets using TRIzol reagent (ThermoFisher Scientific) and cDNA was synthesized as published (17). Gene expression was assessed as previously described (17) using ABI TaqMan commercial primers (Applied Biosystems) except for human *XBP1S* that was assessed using SYBR green master mix (Thermofisher Scientific) using primers in Supplementary Table 3.

### Western Blot

Lysates of INS1 cells and islets were prepared in RIPA buffer as previously described (17). Western blot was performed as described previously (9) using antibodies as listed in Supplementary Table 4. Signal was captured by enhanced chemiluminescence as described (9) except for Figure 9h that utilized Odyssey CLx imaging system (Licor).

### Calcium imaging

INS1 cells were re-plated on a cover slip coated with extracellular matrix of HTB9 cells 72h after transfection and calcium imaging was performed as published (54). In brief, cells were loaded with Fura-2-AM (10 μM) in RPMI1640 at 37 °C for 20 min followed by bathing at 37 °C in the basal KRBH solution containing (in mM): 129 NaCl, 5 NaHCO_3_, 4.8 KCl, 1.2 KH2PO_4_, 2.5 CaCl_2_, 2.4 MgSO_4_, 10 HEPES, 1 glucose, 29 mannitol, 0.1% w/v bovine serum albumin, pH 7.4 with NaOH (300 mOsm/kg) with change to 30 mM glucose or 30 mM KCl at time indicated. INS1 cells were excited by 340 and 387 nm light alternatively using a DG-4 xenon-arc lamp (Sutter Instruments) and emission signals were recorded at 510 nm every 3 s using CMOS charge-coupled device (CCD) camera (Orca flash 4.0+, Hamamatsu). Images of cells were capture by an Olympus IX73 microscope using a 20 x /0.75 NA objective. The ratio of 340/380 fluorescent signal intensity (*I340/I380*) was used as the measure of [Ca^2+^]_i_. Area under the curve (AUC) was obtained from the average tracing for each cover slip. Glucose response was calculated as (AUC during 30 mM glucose-AUC during 1 mM glucose)/AUC during 1 mM glucose.

### Oxygen Consumption Rate

Respirometry of INS1 cells and mouse islets was performed using an Agilent Seahorse XF24 respirometer (Agilent Technologies) as previously described (17). In brief, the OCR was sequentially measured starting from 5 cycles without glucose followed by 5 cycles of 22.2 mM glucose, 6 cycles of 5 μM oligomycin, 5 cycles of 1 μM carbonyl cyanide-*4*-(trifluoromethoxy) phenylhydrazone (FCCP), and 6 cycles of 1 μM rotenone + 0.1μM antimycin A. Glucose response was defined as the difference between the highest OCR during glucose phase and OCR prior to glucose addition. ATP production was defined as the difference between the highest OCR during the glucose phase and OCR at the end of oligomycin phase. Maximal respiration was defined as the difference between the highest OCR during FCCP phase and OCR at the end of assay. Proton leak was defined as the difference between OCR at the end of oligomycin phase and at the end of assay. All data were normalized to DNA content measured by fluorometric assay using Hoechst 33258 (385 nm excitation and 450 nm emission). Percent protein leak was expressed taking basal as 100% (24).

### Rat Mitochondrial DNA

Mitochondrial DNA content was measured by comparing mitochondrial gene (Mt-ND6) level relative to nuclear gene (β actin) by real-time PCR in total DNA isolated from INS1 cells by QIAamp DNA minikit (Qiagen). Rat primers were mt-ND6, 5’-TTGGGGTTGCGGCTATTTAT-3’ (forward) and 5’-ATCCCCGCAAACAATGACCA-3’ (reverse); β-actin, 5’-GCTCTATCAC TGGGCATTGG-3’ (forward) and 5’-CGCAACTCTTAACTCGGAAGA-3’ (reverse) published by Yamazaki et al (55).

### Morphometry of LDs in INS1 cells

INS1 cells were re-plated onto a cover slip as above and incubated in regular INS1 medium containing 8 μM Bodipy C12 (Molecular Probes) at 37 °C at 5% CO_2_ for the last 16 h of culture. After washing 3 times in PBS, cells were fixed for 15 min at 37°C in 4% paraformaldehyde and incubated with 2 μM Bodipy 493/503 (Bodipy 493, Molecular Probes) and 1 μg/ml of DAPI as published (9). Images of the cells were captured by a Zeiss LSM710 microscope and analyzed in a blinded fashion by an independent observer. LDs were first identified in each image using the Bodipy 493 channel as follows. The 12-bit images were converted to 8-bit, gaussian blurred (sigma=1), and a threshold of 25 was used to separate the LD foreground from both background and diffuse cellular staining. After thresholding, the image was converted to a binary mask, adjacent objects were separated using the watershed function, and the analyze particles function was used to record each droplet as a region of interest (ROI, Supplementary File 1). Once each droplet was identified, the original 12-bit image was opened and the Bodipy C12 channel was analyzed at each of the ROI’s that had been identified using the Bodipy 493 channel. For each ROI, the area and integrated intensity was recorded (Supplementary File 2). In addition, the total integrated intensity across the entire image was recorded. The same data extraction was repeated for every image in the dataset. Percent C12 inside LD was calculated by dividing the sum of the integrated intensity of C12 from the ROI’s of every droplet in the image by the integrated intensity of the entire image after background subtraction. INS1 cells labeled with Bodipy C12 were also immunostained with mouse anti-HSP60 antibody (66-41-1-lg, Protein Tech) and Alexa Fluor 488 goat anti-Mouse IgG (A11017, Invitrogen) both at 1:300. Pearson’s coefficients of Bodipy C12 and Alexa Fluor 488 were calculated on single-plane confocal images captured by a Zeiss LSM710 microscope using the Colocalization function in Bitplane Imaris v 9.6 (Oxford Instruments). The lower threshold was set to 20 for both channels and all images.

### Mitochondrial morphology

INS1 cells re-plated onto a cover slip as above were incubated in regular INS1 medium supplemented with 0.4 mM OA at 37 °C at 5% CO_2_ for the last 16 h. MitoTracker Deep Red (ThermoFisher Scientific) was added at final 100 nM and incubation was continued for additional 30 min at 37 °C at 5% CO_2_. Then, cells were fixed for 15 min at 37°C by adding paraformaldehyde directly at final concentration of 4% followed by gentle wash with PBS 3 times. Images of cells were captured by a Zeiss LSM710 microscope and converted to binary images of mitochondrial particles using a custom written NIH ImageJ Marco as published (56, 57). Automated morphometry of mitochondrial particles was further performed using the NIH ImageJ Marco to obtain aspect ratio (major axis/minor axis) and form factor (perimeter^2^/(4π x area)) (56, 57). Fragmented mitochondria was defined as those with aspect ratio below 2.

### [^3^H]OA oxidation

48 h after transfection, 0.5 μCi/0.5 ml of [^3^H]OA (specific activity 24 mCi/mmole, Perkin Elmer) was added to INS1 cells and cultured overnight at 37 °C at 5% CO_2_. The total cellular uptake of [^3^H]OA was measured in cell lysates in RIPA buffer using liquid scintillation counting and corrected for protein contents. Medium was collected and 10% fatty acid free BSA was added to remove unincorporated [^3^H]OA. Then, BSA was precipitated by 0.75N TCA as published (12) and [^3^H]H2O in medium was measured using liquid scintillation counting.

### Metabolomics

INS1 cells transfected with siRNA were analyzed at Metabolomic Core Facility, University of Iowa. After washing with PBS, cells were snap frozen on liquid nitrogen, and stored in −80 °C. After overnight lyophilization, dried cells were extracted into an 1 ml/well of methanol/acetonitrile/water (2:2:1 V/V/V) solution containing 1 ng/μl of D4-Succinate, D8-Valine, D4-Citrate, ^13^C5-Glutamine, ^13^C5-Glutamate, ^13^C6-Lysine, ^13^C5-Methionine, ^13^C3-Serine, and ^13^C11-Tryptophan internal standards. The lyophilized cells were scraped for 25 seconds, transferred to a microcentrifuge tube, and flash frozen in liquid nitrogen. Frozen extracts were thawed in a bath sonicator at room temperature (RT) for 10 min, rotated at −20 °C for 1 h, and centrifuged at 20,000 x g at 4 °C for 10 min. 400 μl of the supernatants transferred into new tubes were dried using a Speedvac for 3 h at RT. Dried extracts were reconstituted in 40 μL acetonitrile/water (1:1 v/v), vortexed for 5 min, and centrifuged at 20,000 x g at 4 °C for 10 min. Finally, 2μL of metabolite extracts were separated using a Millipore SeQuant ZIC-pHILIC (2.1 X 150 mm, 5 μm particle size) column with a ZIC-pHILIC guard column (20 x 2.1 mm) attached to a Thermo Vanquish Flex UHPLC. Mobile phase comprised Buffer A: 20 mM (NH4)^2^CO3, 0.1% NH4OH and Buffer B: acetonitrile. The chromatographic gradient was run at a flow rate of 0.150 mL/min as follows: 0–20 min—linear gradient from 80 to 20% Buffer B; 20–20.5 min—linear gradient from 20 to 80% Buffer B; and 20.5–28 min—hold at 80% Buffer B (58). Data was acquired using a Thermo Q Exactive mass spectrometer operated in full-scan, polarity-switching mode with a spray voltage set to 3.0 kV, the heated capillary held at 275 °C, and the HESI probe held at 350 °C. The sheath gas flow was set to 40 units, the auxiliary gas flow was set to 15 units, and the sweep gas flow was set to 1 unit. MS data was acquired in a range of m/z 70–1,000, with the resolution set at 70,000, the AGC target at 10e6, and the maximum injection time at 200 ms. Acquired LC-MS data were processed by Thermo Scientific TraceFinder 4.1 software, and metabolites were identified based on the University of Iowa Metabolomics Core facility in-house, physical standard-generated library. NOREVA was used for signal drift correction (59). Data per sample were then normalized to total ion signal per sample (Supplementary Table 1) and MetaboAnalyst 4.0 was used for further statistical processing and visualization (60).

### Induction of ER stress

Mouse islets were incubated in 10% FBS RPMI1640 medium with 5 μg/ml of tunicamycin for 6 h. INS1 cells were incubated in INS1 medium with 1 μg/ml of tunicamycin for 6 h. Human islets were incubated in 1% HSA CMRL1066 medium with 10 μg/ml tunicamycin for overnight prior to harvest.

### Human Islets

Human islets from IIDP, Alberta Diabetes Institutes, or Prodo labs (Supplementary Tables 5 and 6) with reported viability and purity above 80% were cultured overnight at 37 °C and 5% CO_2_ upon arrival for recovery from shipping (9). Lentivirus carrying scramble (CCTAAGGTTAAGTCGCCCTCG) and shRNA sequence targeting human PLIN2 (CAGAAGCTAGAGCCGCAAATT) obtained from Genetic Perturbation Platform (https://portals.broadinstitute.org/gpp/public) were prepared as previously described (35). Pseudoislets transduced with lentivirus were created in 96 well spheroid plate (Corning) at 2,000 cells/well or Aggrewell 400 (StemCell technologies) at 400 cells/well as described (61) and cultured for 1 week in CMRL 1066 supplemented with 10% heat inactivated FBS, 1% Pen-Strep, and 1% L-Glutamate (10% HI-FBS CMRL) at 37 °C and 5% CO_2_ before harvesting. The study was determined to be a non-human study by Institutional Review Board at University of Iowa.

### Perifusion of Islets

BioRep Perifusion System (BioRep Technologies) was used to perifuse human pseudoislets as published (35). In brief, after 48 minutes in 2.8 mM glucose in KRB, islets were perifused for indicated time in 16.7 mM glucose or 30 mM KCl in 2.8 mM glucose. Total insulin contents were obtained from islets incubated overnight at 4°C in acidified ethanol. Insulin was measured using STELLUX Chemiluminescent Human Insulin ELISA (ALPCO). Insulin secretion was expressed by taking total insulin contents as 100%. Stimulation index (SI) for the first phase was determined as the average insulin secretion during the first 6 minutes and SI for the second phase as the average of insulin secretion after 6 minutes till the end of glucose ramp, both divided by average basal insulin secretion.

### Statistics

Data are presented as mean ± s.e.m. unless otherwise stated in the figure caption. Differences of numeric parameters between two groups were assessed with Student’s t-tests. Pairwise comparison was used for values obtained in human islets from the same donor. Welch correction was applied when variances between two groups were significantly different by F test using Prism 8 (GraphPad, La Jolla, CA). Multiple group comparisons used one-way ANOVA with post hoc as indicated. A p < 0.05 was considered significant.

## Author Contribution

YI conceived the study and is responsible for all contents of the manuscript. AM (all aspects), SL, (imaging and human islet studies), JP (insulin secretion and mouse studies), MH (human pseudoislet studies), WS, and BF (Seahorse metabolic analyzer), GB and BO (OXPHOS analysis), CK and RS (calcium signaling), SStr (mitochondrial morphology), SSte (INS1 cell studies), TK, and LJ (mouse studies), FA-D and RA (mouse studies), JA (morphometric analysis of LD) were responsible for the acquisition and/or analysis of the data. AG was responsible for establishment of the mouse model and designing of mouse studies. YI, AM, and SL designed research, drafted the manuscript, and critically revised the manuscript for important intellectual content. All authors revised and approved the final version of the manuscript.

## Acknowledgement

This work was financially supported by National Institutes of Health to YI (R01-DK090490). YI, BO, and JA. are supported by the Fraternal Order of Eagles Diabetes Research Center. BO is supported by VA Merit Review Award Number lO1 BXOO4468. AG is supported by ARS Project 1950-51000-071-02S, P30 DK046200-23, P30 DK046200-27S2, DK108722 01A1, 1R21HD098056-01, and support from the Robert C and Veronica Atkins Foundation. Authors utilized human pancreatic islets provided by the NIDDK-funded Integrated Islet Distribution Program (IIDP) at City of Hope (2UC4DK098085). A Zeiss LSM710 confocal microscope (1 S10 RR025439-01) located in the University of Iowa Central Microscopy Research Facility (CMRF) was purchased through the indicated NIH SIG grants. We thank Dr. Thomas Moninger (CMRF Assistant Director) and Drs. Fariba Tayyari and Adam Rauckhorst of the Fraternal Order of Eagles Diabetes Research Center Metabolomic Core, University of Iowa for technical assistance. A part of studies were supported by resources and the use of facilities at the Department of Veterans Affairs Health Care System, Iowa City, IA 52246.

