## Supplemental materials for "Perilipin2 down-regulation in β cells impairs insulin secretion under nutritional stress and damages mitochondria"

**Supplementary File 1. ROI identification ImageJ Macro**

```
run("8-bit");

run("Arrange Channels...", "new=2");

run("Gaussian Blur...", "sigma=1");

setAutoThreshold("Default dark");

//run("Threshold...");

setThreshold(25, 255);

//setThreshold(25, 255);

run("Convert to Mask");

run("Watershed");

run("Analyze Particles...", "size=0.04-Infinity circularity=0.10-1.00 display exclude clear add");
```

**Supplementary File 2. ROI Quantification ImageJ Macro**

```
run("Arrange Channels...", "new=12");

roiManager("multi-measure measure_all");
```

**Methods for supplementary figures**

***Morphometric analysis of beta cell area***

Morphometric analysis of mouse pancreatic sections was performed as published (1). In brief, whole pancreas was dissected en bloc, fixed in 10% formalin overnight, and paraffin embedded to keep the original shape of pancreas. 7 μm section that contains the maximum footprint was deparaffinized, rehydrated, and immunostained with guinea pig anti-insulin antibody (Ab7842, ABCAM at 1:100) followed by the visualization by Alexa 488 anti-guinea pig antibody at 1:500. Islets were imaged using a Leica DMI6000 B Inverted Microscope by first locating one insulin positive area and then adjusting fluorescence intensity for best quality images at 10X magnification. GFP settings used for 10X magnification were 300 ms exposure and gain of 16. Next, the microscope was switched to 5X magnification to allow whole pancreas section imaging. Insulin positive sections were then sequentially located in the 5X magnification and imaged using the 10X magnification using preset settings. Once all insulin positive areas were imaged, size of insulin positive area was quantified automatically using ImageJ analysis software.

***Mouse Mitochondrial DNA***

Mitochondrial DNA content was measured by comparing mitochondrial gene (MTeCO1) level relative to nuclear gene (Ndufv1) by real-time PCR in total DNA isolated from mouse islets by QIAamp DNA minikit (Qiagen). Primers were MTeCO1 [(F) 5’-TGC TAG CCG CAG GCA TTA C-3’ (forward) and 5’-GGG TGC CCA AAG AAT CAG AAC-3’ (Reverse); Ndufv1, 5’-CTT CCC CAC TGG CCT CAA G-3’ (forward) and 5’-CCA AAA CCC AGT GAT CCA GC-3’ (reverse) as published by Doliba et al.(2)

***Principal Component Analysis***

Principal Component Analysis (PCA) was performed using MetaboAnalyst 4.0 software for metabolomic data (Supplementary Table 1) obtained from 2 independent experiments of INS1 cells treated with Scr control and SiPLIN2 as in Methods, each performed in triplicate. A 2D scores plot comparing the top 2 PC calculated is shown in Supplementary Figure 4.

**Supplementary Figure**

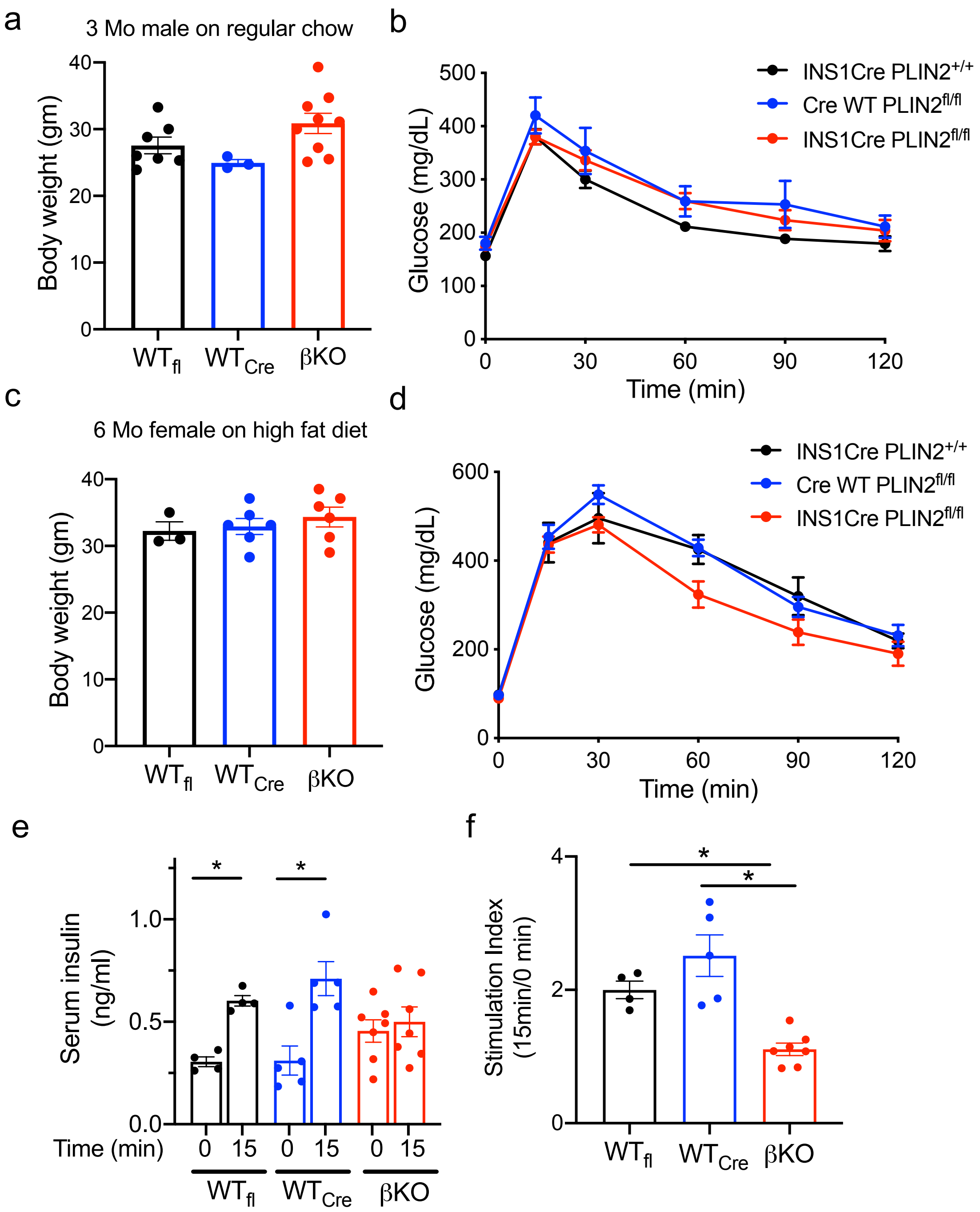


**Figure 1. Metabolic profiling of beta cell specific PLIN2 deficient mice.**
(a) body weight (BW) and (b) 1.5 mg/g BW glucose GTT was done on 3-month old male mice on regular rodent chow after 6 h fasting. $WT_{fl}$; INS1Cre $PLIN2^{+/+}$, $WT_{cre}$; CreWT $PLIN2^{fl/fl}$, βKO; INS1Cre $PLIN2^{fl/fl}$. n= 7 for $WT_{fl}$, 3 for $WT_{cre}$, and 9 for βKO. (c) BW and (d) 1.0 mg/g BW glucose GTT was done on 6-month old female mice on high fat diet chow for 6 weeks after 6 hour

fasting. $WT_{fl}$; INS1Cre $PLIN2^{+/+}$, $WT_{cre}$; CreWT $PLIN2^{fl/fl}$, βKO; INS1Cre $PLIN2^{fl/fl}$. n= 3 for $WT_{fl}$, 6 for $WT_{cre}$, and 6 for βKO. (e) Serum insulin in response to 1.5 mg/g BW i.p. glucose GTT and (f) stimulation index determined as insulin level at 15 min over 0 min for each mouse. n= 4 for $WT_{fl}$, 5 for $WT_{cre}$, and 7 for βKO. All data are mean ± s.e.m.

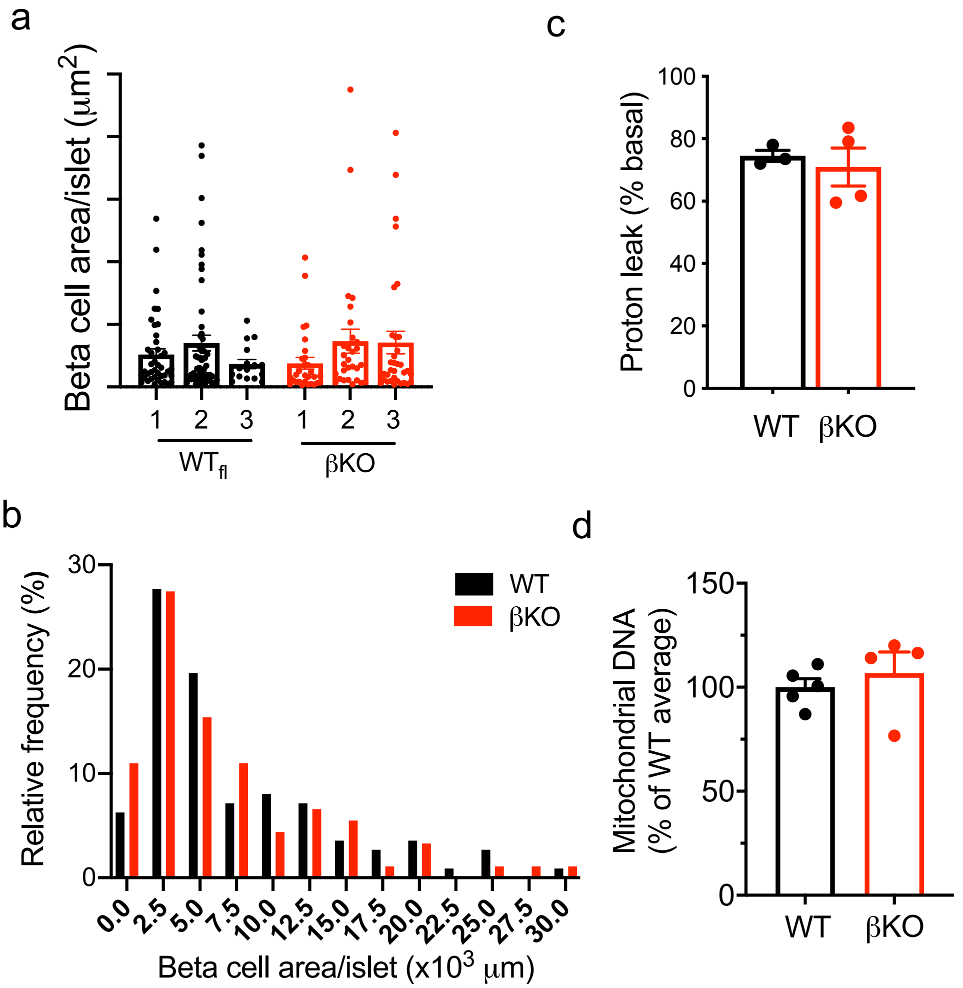


**Figure 2. Islet data from beta cell specific PLIN2 deficient mice.**

(a-b) Beta cell area per islet was defined by measuring insulin positive area as in Supplementary methods in pancreatic section from 3 WT (WT$_{fl}$;INS1Cre PLIN2$^{+/+}$) and 3 βKO (INS1Cre PLIN2$^{fl/fl}$) mice placed on HFD. (a) mean ± s.e.m. of islet size for each mouse. n= 41 for WT1, 55 for WT2, 16 for WT3, 29 for KO1, 29 for KO2, and 33 for KO3. (b) Distribution of islet size defined as beta cell area. Data combines all islets from 3WT and 3 βKO from (a). n=112 for WT and 91 for βKO. (c) Proton leak expressed as % basal oxygen consumption rate (OCR) determined by Seahorse Metabolic analyzer in mouse islets from high fat fed WT (WT$_{fl}$) and βKO islets corrected for DNA. n=4 for WT$_{fl}$ and 3 for βKO. (d) qPCR probed DNA from WT and βKO islets for expression levels of mitochondrial DNA (MTeCO1) and nuclear DNA (Ndufv1). Expression of MTeCO1 was corrected for Ndufv1 and the average of value for WT was taken as 100%. Data are mean ± s.e.m. n=5 for WT (1WT$_{fl}$ and 4WT$_{cre}$) and 4 for βKO.

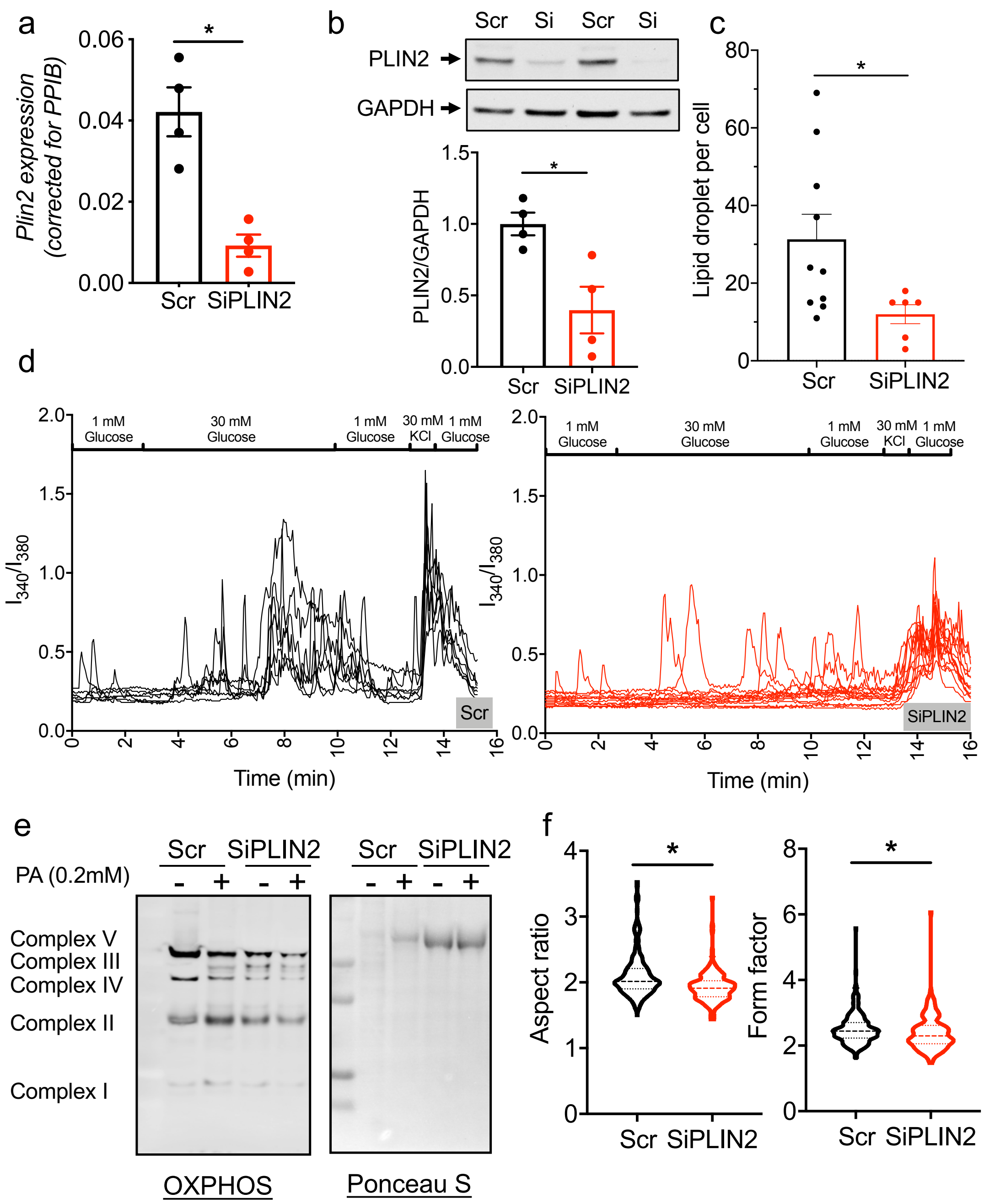


**Figure 3. Down-regulation of PLIN2 in INS1 cells**

(a) Expression of *Plin2* in INS1 cells transfected with scramble control siRNA (Scr) or siRNA targeting Plin2 (SiPlin2) determined by qPCR using *Ppib* as an internal control. n= 4. (b) Western blot compared the expression of PLIN2 in SiPLIN2 and Scr treated INS1 cells. A representative Western blot and densitometry data corrected for GAPDH. n=4. (c) Number of lipid droplets (LDs) per cell in 12 cells treated by Scr and 6 cells treated by SiPLIN2 were measured. Representative of 3 independent experiments. (d) Tracing of Fura-2 $Ca^{2+}$ transients in INS1 cells transfected with Scr or SiPlin2 on a cover slip in response to 30 mM glucose (basal 1 mM glucose) from Fig. 4f plotted to show an individual cell. n= 8 cells for Scr and 14 cells for

SiPLIN2 per cover slide. (e) Representative Western blot for Figure 4h that probed OXPHOS complex proteins in protein lysate of INS1 cells transfected by Scr and SiPLIN2 treated with or without 0.2 mM palmitic acids (PA) for 24 h prior to harvest. (f) Distribution of aspect ratio and form factor shown as violin plots for all mitochondria counted in three independent experiments n= 210 for Scr and 154 for siPLIN2. Medium and quantiles are shown. For all others, data are mean ± s.e.m. *; $p<0.05$ by student's t test.

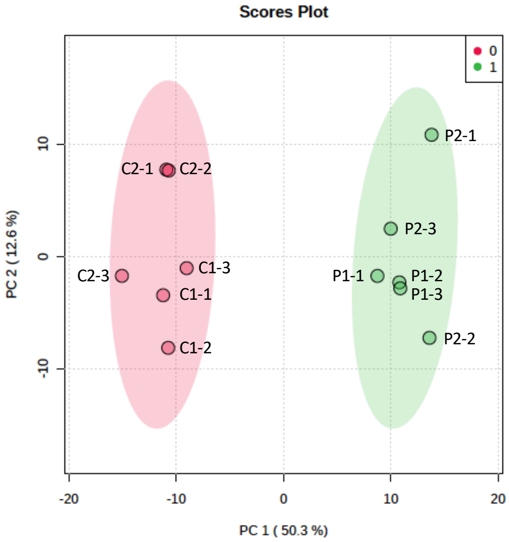


**Figure 4. Principal component analysis compared metabolites between Scr and SiPLIN2 treated INS1 cells**

% variance explained by principal component (PC)1 is shown in x axis and PC2 in y axis. Experiments were repeated twice (Exp1 and 2) each in triplicates and the results were combined making total 6 samples per treatment. Scr control (C1, C2) values were shaded red and siPLIN2 (P1, P2) values in green. See Supplementary Table 1 for raw data.

### Supplementary Table 1: List of metabolites measured

| | | | EXP 1 | | | | | | EXP2 | | | | | |
|---|---|---|---|---|---|---|---|---|---|---|---|---|---|---|
| | | | Scr | | | SiPLIN2 | | | Scr | | | SiPLIN2 | | |
| Name | KEGG[a] | HMDB[b] | 1 | 2 | 3 | 1 | 2 | 3 | 1 | 2 | 3 | 1 | 2 | 3 |
| 2-Deoxy-guanosine 5-triphosphate sodium salt hydrate | C00286 | HMDB 0001440 | 9.39 | 7.75 | 9.67 | 8.94 | 8.48 | 8.26 | 11.77 | 11.05 | 8.83 | 13.53 | 8.20 | 12.60 |
| 4-Imidazoleacrylic acid | C00785 | HMDB 0000301 | 10.80 | 8.68 | 7.02 | 7.44 | 7.87 | 5.30 | 9.31 | 8.89 | 21.25 | 4.57 | 7.36 | 19.17 |
| 5-Deoxy.5. methylthio. adenosine | C00170 | HMDB 0001173 | 8.50 | 7.91 | 8.75 | 11.25 | 9.63 | 9.50 | 9.92 | 9.60 | 10.48 | 10.46 | 10.62 | 10.22 |
| 6-phospho-gluconate | C00345 | HMDB 0001316 | 11.10 | 9.05 | 9.33 | 7.25 | 6.35 | 7.13 | 13.33 | 13.07 | 15.97 | 9.02 | 6.00 | 7.04 |
| Adenine | C00147 | HMDB 0000034 | 8.24 | 8.33 | 8.17 | 11.63 | 9.24 | 9.17 | 10.18 | 10.21 | 10.68 | 11.13 | 10.58 | 11.34 |
| Adenosine | C00212 | HMDB 0000050 | 10.10 | 9.91 | 9.30 | 7.73 | 7.91 | 7.26 | 14.44 | 9.83 | 10.75 | 9.92 | 8.74 | 8.53 |
| ADP | C00008 | HMDB 0001341 | 9.01 | 5.35 | 8.49 | 7.97 | 5.82 | 7.24 | 9.29 | 12.18 | 7.84 | 14.07 | 12.81 | 11.40 |
| Alanine | C00041 | HMDB 0000161 | 8.89 | 8.94 | 9.10 | 7.99 | 8.35 | 7.70 | 11.80 | 11.70 | 12.04 | 9.12 | 9.36 | 8.66 |
| Allantoin | C01551 | HMDB 0000462 | 9.21 | 9.25 | 8.52 | 8.76 | 8.62 | 8.38 | 10.37 | 10.91 | 10.85 | 10.63 | 11.07 | 9.66 |
| AMP | C00020 | HMDB 0000045 | 9.33 | 9.14 | 9.39 | 8.05 | 8.54 | 8.31 | 11.28 | 11.33 | 9.03 | 11.41 | 10.25 | 11.51 |
| ATP | C00002 | HMDB 0000538 | 9.39 | 7.75 | 9.67 | 8.94 | 8.48 | 8.26 | 11.77 | 11.05 | 8.83 | 13.53 | 8.20 | 12.60 |
| B.Nicotinamide. mononucleotide | C00455 | HMDB 0000229 | 4.04 | 5.18 | 5.91 | 8.72 | 11.08 | 10.40 | 8.19 | 12.15 | 9.93 | 10.79 | 12.08 | 9.20 |
| Butyryl.L.carnitine | C02862 | HMDB 0002013 | 8.96 | 9.65 | 8.85 | 8.92 | 8.27 | 8.84 | 11.04 | 10.99 | 10.37 | 9.89 | 9.49 | 9.36 |
| Citrulline | C00327 | HMDB 0000904 | 7.64 | 8.37 | 8.38 | 8.84 | 9.82 | 9.18 | 10.00 | 9.81 | 10.03 | 11.02 | 11.33 | 11.22 |
| CMP | C00055 | HMDB 0000095 | 11.78 | 10.46 | 15.07 | 7.60 | 9.62 | 7.81 | 12.01 | 12.27 | 11.77 | 9.68 | 9.07 | 7.94 |
| Creatine | C00300 | HMDB 0000064 | 9.12 | 9.30 | 8.86 | 8.34 | 8.79 | 7.39 | 9.86 | 11.05 | 10.80 | 9.72 | 10.47 | 9.52 |
| Cytidine | C00475 | HMDB 0000089 | 9.93 | 10.04 | 9.82 | 7.98 | 8.69 | 7.80 | 12.82 | 12.31 | 13.67 | 7.91 | 7.45 | 7.75 |
| Cytidine.5.diphosphocholine.sodium.salt. dihydrate | C00307 | HMDB 0001413 | 7.98 | 9.19 | 9.26 | 7.20 | 9.62 | 8.01 | 10.09 | 9.90 | 9.43 | 10.45 | 12.24 | 10.76 |
| Decanoyl.L.carnitine | C03299 | HMDB 0062631 | 9.04 | 13.38 | 13.48 | 12.21 | 13.41 | 12.81 | 7.59 | 9.48 | 10.49 | 12.61 | 11.39 | 11.79 |
| D.Gluconic.acid.solution | C00257 | HMDB 0000625 | 8.66 | 8.61 | 8.50 | 9.14 | 8.75 | 9.05 | 11.31 | 11.79 | 11.49 | 10.16 | 10.38 | 9.98 |
| D.Glucose.6.phosphate.potassium.salt | C00092 | HMDB 0001401 | 10.43 | 9.89 | 9.78 | 7.46 | 7.68 | 7.43 | 11.81 | 11.89 | 10.81 | 9.57 | 9.16 | 9.60 |
| DL. Homocysteine | C05330 | HMDB 0000742 | 9.32 | 7.65 | 9.93 | 7.97 | 8.54 | 8.69 | 11.01 | 11.41 | 9.01 | 10.58 | 9.52 | 11.27 |
| DL. lauroylcarnitine | None | HMDB 0002250 | 12.24 | 12.13 | 9.48 | 10.86 | 11.13 | 14.98 | 8.20 | 9.20 | 9.98 | 10.96 | 12.38 | 12.11 |
| DL.Leucine | C00123 | HMDB 0000687 | 5.18 | 7.29 | 7.91 | 9.20 | 10.42 | 10.65 | 11.24 | 10.12 | 10.89 | 11.81 | 11.89 | 10.75 |
| D.Ornithine.monohydrochloride | C00515 | HMDB 0003374 | 9.10 | 8.48 | 8.40 | 8.41 | 8.78 | 8.78 | 10.45 | 10.59 | 9.66 | 10.22 | 10.08 | 9.95 |
| D.Tryptophan | C00525 | HMDB 0013609 | 8.09 | 7.90 | 7.55 | 10.74 | 9.95 | 10.32 | 9.89 | 9.67 | 9.74 | 11.72 | 11.63 | 10.68 |
| FAD | C00016 | HMDB 0001248 | 9.47 | 8.73 | 9.48 | 8.35 | 8.66 | 8.45 | 10.49 | 10.41 | 8.82 | 11.47 | 9.60 | 10.52 |

| | | | | | | | | | | | | | | |
|---|---|---|---|---|---|---|---|---|---|---|---|---|---|---|
| GDP | C00035 | HMDB 0001201 | 8.72 | 8.96 | 8.95 | 8.62 | 9.65 | 7.87 | 11.12 | 13.61 | 9.10 | 14.61 | 9.51 | 12.01 |
| Glutamate | C00025 | HMDB 0000148 | 8.54 | 8.90 | 8.49 | 8.95 | 9.32 | 8.85 | 10.84 | 11.33 | 11.14 | 9.49 | 9.49 | 9.24 |
| Glutamine | C00064 | HMDB 0000641 | 5.99 | 5.94 | 5.73 | 12.1 6 | 11.65 | 11.53 | 9.74 | 9.09 | 9.69 | 11.33 | 11.47 | 10.78 |
| Glutathione. oxidized. | C00127 | HMDB 0003337 | 7.53 | 8.05 | 7.56 | 8.81 | 9.11 | 8.80 | 8.86 | 8.27 | 8.43 | 12.11 | 10.79 | 12.31 |
| Glutathione. reduced. | C00051 | HMDB 0000125 | 9.69 | 8.95 | 8.28 | 9.89 | 8.61 | 9.14 | 8.94 | 9.26 | 9.86 | 11.42 | 11.86 | 11.35 |
| Glycerol | C00116 | HMDB 0000131 | 10.60 | 8.38 | 9.01 | 9.47 | 7.53 | 7.57 | 8.20 | 7.96 | 7.51 | 7.07 | 20.30 | 6.68 |
| Glycine | C00037 | HMDB 0000123 | 9.02 | 9.33 | 9.21 | 8.35 | 8.93 | 8.94 | 11.92 | 11.33 | 11.34 | 8.93 | 8.74 | 9.00 |
| GMP | C00144 | HMDB 0001397 | 9.77 | 9.29 | 9.76 | 7.61 | 9.14 | 8.55 | 8.80 | 10.43 | 9.23 | 12.27 | 11.32 | 11.45 |
| GTP | C00044 | HMDB 0001273 | 8.34 | 7.04 | 8.69 | 9.27 | 7.92 | 7.81 | 14.29 | 16.82 | 9.42 | 18.03 | 6.87 | 12.62 |
| Guanosine | C00387 | HMDB 0000133 | 9.65 | 9.94 | 9.01 | 7.99 | 8.51 | 7.90 | 10.27 | 10.52 | 9.85 | 10.22 | 9.80 | 9.75 |
| Hexanoyl. L.carnitine | None | HMDB 0000756 | 8.98 | 8.80 | 8.38 | 8.69 | 8.61 | 8.13 | 8.93 | 11.41 | 11.00 | 10.98 | 10.52 | 11.00 |
| Hypotaurine | C00519 | HMDB 0000965 | 8.12 | 8.11 | 7.76 | 9.23 | 9.53 | 9.01 | 10.42 | 10.40 | 10.84 | 10.07 | 10.62 | 10.48 |
| Hypoxanthine | C00262 | HMDB 0000157 | 7.42 | 6.92 | 6.26 | 9.69 | 10.85 | 10.91 | 5.60 | 5.71 | 5.93 | 14.84 | 16.56 | 14.84 |
| IMP | C00130 | HMDB 0000175 | 8.04 | 9.50 | 9.21 | 9.49 | 11.26 | 10.68 | 7.20 | 7.50 | 6.82 | 14.44 | 13.78 | 12.74 |
| Inosine | C00294 | HMDB 0000195 | 8.93 | 9.76 | 8.96 | 9.80 | 10.67 | 11.96 | 8.79 | 9.11 | 10.03 | 10.25 | 11.87 | 10.61 |
| Isoleucine | C00407 | HMDB 0000172 | 9.74 | 9.58 | 10.79 | 9.46 | 8.81 | 9.78 | 10.10 | 9.88 | 10.66 | 11.11 | 11.26 | 10.70 |
| L. Acetylcarnitine | C02571 | HMDB 0000201 | 10.03 | 11.05 | 9.27 | 7.59 | 6.90 | 7.39 | 11.73 | 13.56 | 13.07 | 7.81 | 7.63 | 7.52 |
| Lactate | C00186 | HMDB 0000190 | 9.77 | 8.65 | 9.12 | 8.20 | 8.11 | 8.50 | 10.88 | 10.74 | 11.11 | 9.96 | 9.52 | 9.44 |
| L.Asparagine | C00152 | HMDB 0000168 | 8.96 | 8.82 | 8.84 | 9.00 | 9.15 | 8.49 | 11.02 | 10.63 | 10.87 | 9.73 | 10.18 | 9.73 |
| L.Aspartic.acid | C00049 | HMDB 0000191 | 7.61 | 7.61 | 7.43 | 10.4 2 | 9.95 | 9.89 | 10.50 | 10.05 | 10.49 | 10.58 | 10.47 | 10.17 |
| L.Carnitine | C00318 | HMDB 0000062 | 10.00 | 10.61 | 8.55 | 8.41 | 7.41 | 8.05 | 9.98 | 12.28 | 10.99 | 9.26 | 9.44 | 9.14 |
| L.Carnosine | C00386 | HMDB 0000033 | 7.83 | 7.06 | 8.88 | 9.76 | 11.14 | 10.26 | 10.24 | 9.36 | 9.35 | 10.60 | 11.20 | 10.91 |
| L.Cystathionine | C02291 | HMDB 0000099 | 8.65 | 9.80 | 8.89 | 9.70 | 9.80 | 8.86 | 9.67 | 12.03 | 10.10 | 9.88 | 11.34 | 10.41 |
| L.Dihydroorotic. acid | C00337 | HMDB 0003349 | 8.21 | 5.60 | 6.18 | 8.66 | 6.32 | 8.30 | 7.41 | 10.81 | 10.59 | 10.64 | 12.13 | 10.53 |
| L.Homoserine | C00263 | HMDB 0000719 | 7.87 | 8.06 | 8.11 | 8.51 | 9.26 | 8.22 | 11.05 | 9.46 | 10.57 | 10.30 | 10.36 | 9.03 |
| L.Kynurenine | C00328 | HMDB 0000684 | 8.24 | 8.19 | 9.08 | 8.25 | 10.36 | 13.25 | 11.96 | 11.07 | 11.54 | 11.77 | 12.05 | 10.06 |
| L.Malic.acid | C00149 | HMDB 0000156 | 9.76 | 9.78 | 9.11 | 7.92 | 7.50 | 7.71 | 11.27 | 12.04 | 11.39 | 8.70 | 9.00 | 8.89 |
| L.Phenylalanine | C00079 | HMDB 0000159 | 7.37 | 7.43 | 7.54 | 10.1 1 | 10.69 | 10.59 | 10.76 | 9.53 | 9.68 | 11.22 | 11.52 | 10.58 |
| L.Tartaric.acid | C00898 | HMDB 00956 | 7.57 | 8.43 | 12.59 | 6.18 | 7.39 | 10.93 | 8.30 | 11.33 | 13.71 | 9.53 | 6.27 | 11.19 |
| L.Tyrosine | C00082 | HMDB 0000158 | 7.06 | 7.18 | 7.56 | 9.56 | 10.40 | 10.43 | 11.02 | 10.03 | 9.56 | 10.79 | 11.10 | 11.02 |
| L.Valine | C00183 | HMDB 0000883 | 8.90 | 8.23 | 8.23 | 9.20 | 9.45 | 8.97 | 11.29 | 10.46 | 11.62 | 10.69 | 10.88 | 10.52 |
| Malonic.acid | C00383 | HMDB 0000691 | 6.09 | 7.13 | 10.20 | 6.06 | 6.74 | 10.92 | 10.38 | 10.16 | 10.48 | 10.37 | 11.02 | 12.57 |
| Methionine | C00073 | HMDB 0000696 | 7.76 | 7.44 | 6.80 | 9.34 | 9.79 | 10.68 | 9.91 | 8.52 | 9.72 | 10.67 | 12.70 | 12.39 |
| N.Acetyl. L.aspartic.acid | C01042 | HMDB 0000812 | 7.23 | 8.29 | 6.82 | 9.66 | 10.16 | 10.24 | 8.05 | 8.51 | 7.65 | 13.79 | 12.30 | 12.72 |
| N.acetyl. neuraminic.acid | C19910 | HMDB 0000230 | 11.73 | 10.84 | 10.64 | 6.05 | 6.51 | 6.66 | 13.94 | 14.08 | 13.43 | 7.48 | 7.17 | 7.53 |

| | | | | | | | | | | | | | | |
|---|---|---|---|---|---|---|---|---|---|---|---|---|---|---|
| NAD | C00003 | HMDB 0000902 | 9.19 | 9.62 | 9.08 | 8.81 | 9.09 | 8.37 | 12.04 | 11.55 | 10.76 | 10.42 | 9.90 | 9.66 |
| NADH | C00004 | HMDB 0001487 | 9.53 | 7.42 | 10.91 | 11.33 | 7.84 | 10.49 | 14.25 | 9.38 | 8.89 | 12.78 | 8.59 | 12.71 |
| NADP | C00006 | HMDB 0000217 | 9.49 | 7.85 | 9.75 | 7.62 | 7.90 | 8.29 | 9.34 | 11.61 | 8.67 | 12.76 | 9.76 | 10.90 |
| NADPH | C00005 | HMDB 00221 | 8.57 | 7.02 | 8.26 | 9.56 | 9.25 | 8.42 | 13.32 | 8.26 | 8.95 | 12.25 | 8.73 | 12.44 |
| N.alpha.Acetyl.L. asparagine | None | HMDB 0006028 | 7.63 | 8.94 | 8.74 | 10.59 | 10.43 | 8.81 | 10.45 | 10.29 | 10.93 | 10.32 | 11.29 | 9.64 |
| Nicotinamide | C00153 | HMDB 0001406 | 9.03 | 8.90 | 9.10 | 9.11 | 7.49 | 8.13 | 10.80 | 10.21 | 11.55 | 10.38 | 11.92 | 10.58 |
| Nicotinamide. hypoxanthine. dinucleotide. sodium.salt | C04423 | None | 7.23 | 7.15 | 8.93 | 9.87 | 12.45 | 10.45 | 9.65 | 8.97 | 6.40 | 10.62 | 13.38 | 11.65 |
| Nicotinate | C00253 | HMDB 0001488 | 9.00 | 8.22 | 10.15 | 8.06 | 8.69 | 8.64 | 11.56 | 9.09 | 10.68 | 10.05 | 10.85 | 10.14 |
| N.Methyl. D.aspartic.acid | C12269 | HMDB 0002393 | 8.07 | 8.32 | 8.11 | 9.68 | 10.17 | 9.56 | 10.74 | 10.87 | 10.53 | 9.72 | 9.85 | 9.53 |
| Octanoyl-L-carnitine | C02838 | HMDB 0000791 | 9.41 | 7.43 | 13.65 | 9.62 | 8.08 | 10.73 | nd | nd | nd | nd | nd | nd |
| Oleoyl. L.carnitine | | HMDB 0005065 | 11.85 | 12.80 | 10.14 | 9.95 | 8.89 | 9.69 | 8.97 | 9.26 | 11.16 | 9.75 | 12.17 | 12.35 |
| O.Phosphorylethanolamine | C00346 | HMDB 0000224 | 9.42 | 8.16 | 8.18 | 8.68 | 8.71 | 8.61 | 8.31 | 8.48 | 8.46 | 12.07 | 12.22 | 12.58 |
| Palmitoyl. L.carnitine | C02990 | HMDB 0000222 | 9.50 | 11.24 | 8.37 | 7.91 | 6.09 | 7.37 | 10.22 | 10.28 | 13.09 | 8.52 | 10.15 | 10.63 |
| Phosphocreatine | C02305 | HMDB 0001511 | 8.81 | 8.53 | 8.93 | 8.38 | 8.96 | 8.26 | 11.00 | 10.24 | 10.11 | 10.12 | 11.30 | 10.07 |
| Proline | C00148 | HMDB 0000162 | 8.04 | 8.26 | 8.37 | 8.77 | 9.28 | 8.37 | 11.52 | 11.23 | 10.80 | 9.90 | 10.29 | 9.95 |
| Propionyl. L.carnitine | C03017 | HMDB 0062514 | 10.06 | 11.67 | 9.16 | 8.11 | 7.29 | 7.65 | 10.89 | 12.51 | 12.46 | 0.07 | 8.26 | 7.64 |
| Pyridoxine | C00314 | HMDB 0000239 | 10.36 | 11.26 | 8.34 | 7.49 | 6.31 | 8.30 | 12.21 | 10.04 | 10.72 | 10.24 | 10.68 | 9.28 |
| Ribose.5. phosphate | C00117 | HMDB 0001548 | 12.19 | 10.82 | 9.73 | 7.70 | 6.23 | 7.49 | 14.42 | 14.45 | 16.50 | 5.74 | 5.96 | 6.05 |
| S.5.Adenosyl. L.methionine.p. toluenesulfonate. salt | C00019 | HMDB 0001185 | 6.95 | 6.91 | 7.51 | 10.18 | 11.60 | 10.69 | 8.44 | 8.20 | 8.50 | 11.99 | 12.06 | 11.82 |
| Serine | C00065 | HMDB 0000187 | 10.44 | 10.61 | 10.31 | 7.50 | 6.97 | 7.32 | 12.47 | 12.58 | 11.65 | 8.05 | 8.41 | 8.45 |
| Stearoyl. L.carnitine | None | HMDB 0000848 | 13.01 | 13.40 | 12.74 | 8.03 | 6.41 | 5.22 | 10.47 | 9.96 | 14.65 | 7.63 | 9.00 | 9.09 |
| Succinic.acid | C00042 | HMDB 0000254 | 9.41 | 10.64 | 11.71 | 6.52 | 7.93 | 7.37 | 13.49 | 12.04 | 12.32 | 8.34 | 8.46 | 9.92 |
| Taurine | C00245 | HMDB 0000251 | 8.75 | 8.77 | 8.25 | 9.14 | 8.94 | 8.85 | 9.97 | 9.93 | 10.58 | 10.37 | 10.54 | 10.28 |
| Threonine | C00188 | HMDB 0000167 | 8.27 | 9.34 | 7.72 | 10.13 | 9.46 | 8.30 | 11.39 | 10.61 | 10.64 | 9.73 | 9.79 | 8.71 |
| trans.4.Hydroxy. L.proline | C01157 | HMDB 0000725 | 8.15 | 8.45 | 8.21 | 9.27 | 8.93 | 8.40 | 11.10 | 9.70 | 11.16 | 9.93 | 10.47 | 9.78 |
| UMP | C00105 | HMDB 0000288 | 9.24 | 8.89 | 8.92 | 9.25 | 8.92 | 7.84 | 10.60 | 10.96 | 9.47 | 12.29 | 11.04 | 10.56 |
| Uracil | C00106 | HMDB 0000300 | 8.68 | 9.26 | 8.49 | 9.21 | 9.92 | 10.01 | 10.18 | 9.90 | 10.38 | 11.00 | 11.40 | 10.54 |
| Urate | C00366 | HMDB 0000289 | 9.45 | 8.52 | 8.35 | 8.34 | 8.55 | 8.89 | 11.16 | 9.68 | 11.42 | 10.04 | 11.35 | 8.48 |
| Uridine | C00299 | HMDB 0000296 | 10.63 | 11.25 | 10.13 | 8.90 | 9.37 | 7.95 | 13.55 | 13.05 | 12.65 | 8.60 | 8.17 | 8.19 |
| Valeryl. L.carnitine | None | HMDB 0013128 | 5.77 | 6.09 | 5.56 | 13.34 | 12.14 | 11.11 | 5.10 | 5.60 | 5.31 | 15.55 | 15.65 | 16.79 |
| Xanthine | C00385 | HMDB 0000292 | 8.95 | 9.17 | 7.64 | 9.17 | 8.84 | 9.80 | 8.76 | 9.06 | 9.83 | 12.14 | 12.54 | 11.26 |

a; Compound entry in Kyoto Encyclopedia of Genes and Genomes
b; The Human Metabolome Database
nd; not done

### Supplementary Table 2. Metabolites with significant differences

| | Exp 1 | | | | | | Exp 2 | | | | | | | |
|---|---|---|---|---|---|---|---|---|---|---|---|---|---|---|
| | Scr | | | SiPLIN2 | | | Scr | | | SiPLIN2 | | | Fold[a] change | P value[b] |
| Name | 1 | 2 | 3 | 1 | 2 | 3 | 1 | 2 | 3 | 1 | 2 | 3 | | |
| S.5.Adenosyl.L.methionine.p.toluenesulfonate.salt | 0.98 | 0.97 | 1.05 | 1.43 | 1.63 | 1.50 | 1.01 | 0.98 | 1.01 | 1.43 | 1.44 | 1.41 | 1.47 | 1.20E-07 |
| N.Acetyl.L.aspartic.acid | 0.97 | 1.11 | 0.92 | 1.30 | 1.36 | 1.37 | 1.00 | 1.05 | 0.95 | 1.71 | 1.52 | 1.58 | 1.47 | 4.93E-05 |
| Methionine | 1.06 | 1.01 | 0.93 | 1.27 | 1.34 | 1.46 | 1.06 | 0.91 | 1.04 | 1.14 | 1.35 | 1.32 | 1.31 | 1.07E-04 |
| Citrulline | 0.94 | 1.03 | 1.03 | 1.09 | 1.21 | 1.13 | 1.01 | 0.99 | 1.01 | 1.11 | 1.14 | 1.13 | 1.13 | 1.10E-04 |
| Valeryl.L.carnitine | 0.99 | 1.05 | 0.96 | 2.30 | 2.09 | 1.91 | 0.96 | 1.05 | 1.00 | 2.91 | 2.93 | 3.15 | 2.55 | 6.75E-04 |
| Nicotinamide.hypoxanthine.dinucleotide.sodium.salt | 0.93 | 0.92 | 1.15 | 1.27 | 1.60 | 1.34 | 1.16 | 1.08 | 0.77 | 1.27 | 1.61 | 1.40 | 1.42 | 8.52E-04 |
| D.Tryptophan | 1.03 | 1.01 | 0.96 | 1.37 | 1.27 | 1.32 | 1.01 | 0.99 | 1.00 | 1.20 | 1.19 | 1.09 | 1.24 | 1.54E-03 |
| Uracil | 0.99 | 1.05 | 0.96 | 1.04 | 1.13 | 1.14 | 1.00 | 0.97 | 1.02 | 1.08 | 1.12 | 1.04 | 1.09 | 1.93E-03 |
| L.Carnosine | 0.99 | 0.89 | 1.12 | 1.23 | 1.41 | 1.30 | 1.06 | 0.97 | 0.97 | 1.10 | 1.16 | 1.13 | 1.22 | 3.24E-03 |
| Glutathione.oxidized. | 0.98 | 1.04 | 0.98 | 1.14 | 1.18 | 1.14 | 1.04 | 0.97 | 0.99 | 1.42 | 1.27 | 1.45 | 1.27 | 4.49E-03 |
| Hypoxanthine | 1.08 | 1.01 | 0.91 | 1.41 | 1.58 | 1.59 | 0.97 | 0.99 | 1.03 | 2.58 | 2.88 | 2.58 | 2.10 | 8.42E-03 |
| Xanthine | 1.04 | 1.07 | 0.89 | 1.07 | 1.03 | 1.14 | 0.95 | 0.98 | 1.07 | 1.32 | 1.36 | 1.22 | 1.19 | 1.21E-02 |
| L.Tyrosine | 0.97 | 0.99 | 1.04 | 1.31 | 1.43 | 1.43 | 1.08 | 0.98 | 0.94 | 1.06 | 1.09 | 1.08 | 1.23 | 2.30E-02 |
| IMP | 0.90 | 1.07 | 1.03 | 1.06 | 1.26 | 1.20 | 1.00 | 1.05 | 0.95 | 2.01 | 1.92 | 1.78 | 1.54 | 2.31E-02 |
| Glutamine | 1.02 | 1.01 | 0.97 | 2.07 | 1.98 | 1.96 | 1.02 | 0.96 | 1.02 | 1.19 | 1.21 | 1.13 | 1.59 | 2.44E-02 |
| DL.Leucine | 0.76 | 1.07 | 1.16 | 1.35 | 1.53 | 1.57 | 1.05 | 0.94 | 1.01 | 1.10 | 1.11 | 1.00 | 1.28 | 3.52E-02 |
| Glutathione.reduced. | 1.08 | 1.00 | 0.92 | 1.10 | 0.96 | 1.02 | 0.96 | 0.99 | 1.05 | 1.22 | 1.27 | 1.21 | 1.13 | 4.23E-02 |
| DL.lauroylcarnitine | 1.08 | 1.08 | 0.84 | 0.96 | 0.99 | 1.33 | 0.90 | 1.01 | 1.09 | 1.20 | 1.36 | 1.33 | 1.19 | 4.59E-02 |
| 5-Deoxy.5.methylthio.adenosine | 1.01 | 0.94 | 1.04 | 1.34 | 1.15 | 1.13 | 0.99 | 0.96 | 1.05 | 1.05 | 1.06 | 1.02 | 1.13 | 4.66E-02 |
| N.acetyl.neuraminic.acid | 1.06 | 0.98 | 0.96 | 0.55 | 0.59 | 0.60 | 1.01 | 1.02 | 0.97 | 0.54 | 0.52 | 0.55 | 0.56 | 6.73E-10 |
| Serine | 1.00 | 1.02 | 0.99 | 0.72 | 0.67 | 0.70 | 1.02 | 1.03 | 0.95 | 0.66 | 0.69 | 0.69 | 0.69 | 9.52E-10 |
| L.Malic.acid | 1.02 | 1.02 | 0.95 | 0.83 | 0.79 | 0.81 | 0.97 | 1.04 | 0.98 | 0.75 | 0.78 | 0.77 | 0.79 | 3.03E-07 |
| D.Glucose.6.phosphate.potassium.salt | 1.04 | 0.99 | 0.97 | 0.74 | 0.77 | 0.74 | 1.03 | 1.03 | 0.94 | 0.83 | 0.80 | 0.83 | 0.79 | 3.85E-06 |
| L.Acetylcarnitine | 0.99 | 1.09 | 0.92 | 0.75 | 0.68 | 0.73 | 0.92 | 1.06 | 1.02 | 0.61 | 0.60 | 0.59 | 0.66 | 9.54E-06 |
| Succinic.acid | 0.89 | 1.01 | 1.11 | 0.62 | 0.75 | 0.70 | 1.07 | 0.95 | 0.98 | 0.66 | 0.67 | 0.79 | 0.70 | 2.26E-05 |
| Ribose.5.phosphate | 1.12 | 0.99 | 0.89 | 0.71 | 0.57 | 0.69 | 0.95 | 0.96 | 1.09 | 0.38 | 0.39 | 0.40 | 0.52 | 5.40E-05 |
| 6-phosphogluconate | 1.13 | 0.92 | 0.95 | 0.74 | 0.65 | 0.73 | 0.94 | 0.93 | 1.13 | 0.64 | 0.42 | 0.50 | 0.61 | 1.52E-04 |
| Lactate | 1.06 | 0.94 | 0.99 | 0.89 | 0.88 | 0.93 | 1.00 | 0.98 | 1.02 | 0.91 | 0.87 | 0.87 | 0.89 | 2.10E-04 |
| CMP | 0.95 | 0.84 | 1.21 | 0.61 | 0.77 | 0.63 | 1.00 | 1.02 | 0.98 | 0.81 | 0.75 | 0.66 | 0.71 | 6.15E-04 |
| Uridine | 1.00 | 1.05 | 0.95 | 0.83 | 0.88 | 0.75 | 1.04 | 1.00 | 0.97 | 0.66 | 0.62 | 0.63 | 0.73 | 1.04E-03 |
| Stearoyl.L.carnitine | 1.00 | 1.03 | 0.98 | 0.62 | 0.49 | 0.40 | 0.90 | 0.85 | 1.25 | 0.65 | 0.77 | 0.78 | 0.62 | 1.05E-03 |
| Cytidine | 1.00 | 1.01 | 0.99 | 0.80 | 0.87 | 0.79 | 0.99 | 0.95 | 1.06 | 0.61 | 0.58 | 0.60 | 0.71 | 1.93E-03 |

| | | | | | | | | | | | | | | |
|---|---|---|---|---|---|---|---|---|---|---|---|---|---|---|
| L.Carnitine | 1.03 | 1.09 | 0.88 | 0.87 | 0.76 | 0.83 | 0.90 | 1.11 | 0.99 | 0.84 | 0.85 | 0.82 | 0.83 | 1.99E-03 |
| Alanine | 0.99 | 1.00 | 1.01 | 0.89 | 0.93 | 0.86 | 1.00 | 0.99 | 1.02 | 0.77 | 0.79 | 0.73 | 0.83 | 2.53E-03 |
| NAD | 0.99 | 1.03 | 0.98 | 0.95 | 0.98 | 0.90 | 1.05 | 1.01 | 0.94 | 0.91 | 0.86 | 0.84 | 0.91 | 5.35E-03 |
| Butyryl. L.carnitine | 0.98 | 1.05 | 0.97 | 0.98 | 0.90 | 0.97 | 1.02 | 1.02 | 0.96 | 0.92 | 0.88 | 0.87 | 0.92 | 6.06E-03 |
| Adenosine | 1.03 | 1.01 | 0.95 | 0.79 | 0.81 | 0.74 | 1.24 | 0.84 | 0.92 | 0.85 | 0.75 | 0.73 | 0.78 | 8.50E-03 |
| Propionyl. L.carnitine | 0.98 | 1.13 | 0.89 | 0.79 | 0.71 | 0.74 | 0.91 | 1.05 | 1.04 | 0.01 | 0.69 | 0.64 | 0.60 | 9.16E-03 |
| Glycine | 0.98 | 1.02 | 1.00 | 0.91 | 0.97 | 0.97 | 1.03 | 0.98 | 0.98 | 0.77 | 0.76 | 0.78 | 0.86 | 1.98E-02 |
| Palmitoyl. L.carnitine | 0.98 | 1.16 | 0.86 | 0.82 | 0.63 | 0.76 | 0.91 | 0.92 | 1.17 | 0.76 | 0.91 | 0.95 | 0.80 | 2.06E-02 |
| Creatine | 1.00 | 1.02 | 0.97 | 0.92 | 0.97 | 0.81 | 0.93 | 1.05 | 1.02 | 0.92 | 0.99 | 0.90 | 0.92 | 2.15E-02 |

Values are converted from supplementary table 1 taking the average of three scr as 1 for each experiment
a; Fold change= (Average of SiPLIN2 from Exp 1 and Exp 2)/ (Average of Scr from Exp 1 and Exp 2)
b; p value of student's t test comparing 6 sample of SiPLIN2 from Exp 1 and Exp 2 vs, 6 samples Scr from Exp 1 and Exp 2

**Supplementary Table 3. Human primers used for Sybr green qPCR**

| | FORWARD | REVERSE |
|---|---|---|
| ACTB[ref3] | GAAGATCAAGATCATTGCTCCT | TACTCCTGCTTGCTGATCCA |
| HPRT1[ref3] | CCTGGCGTCGTGATTAGTGAT | AGACGTTCAGTCCTGTCCATAA |
| XBP1S [ref4] | TGCTGAGTCCGCAGCAGGTG | GCTGGCAGGCTCTGGGGAAG |

ACTB; Actin, beta, HPRT1; Hypoxanthinephosphoribosyl transferase1, XBP1s; X-box binding protein, spliced form

**Supplementary Table 4. Antibodies used for Western blots**

| | | | |
|---|---|---|---|
| guinea pig anti-ADFP | 1:5000 | ProSci[a] | Figure 1b |
| rabbit anti-GAPDH (14C10) | 1:1000 | 2118, Cell Signaling Technology | Figure 1b, 2h, 8e, 8g |
| rodent anti-Total OXPHOS | 1:1,000 | ab110413, abcam | Figure 2h, 4h, 9h and supplementary figure 3e |
| Rabbit anti-ADFP | 1:1,000 | Ab52335, abcam | Figure 8e, 8g and supplementary 3b |
| Mouse anti-tubulin (TU-02) | 1:3,000 | sc-8035, Santa Cruz Biotechnology | Figure 8e, 8g, 9h |
| HRP-conjugated goat anti-guinea pig IgG | 1:10,000 | sc-2438, Santa Cruz Biotechnology | Figure 1b |
| HRP-conjugated mouse IgG kappa binding protein (m-IgGκ BP)-IgG | 1:3,000 | sc-516102, Santa Cruz Biotechnology | Figure 8e, 8g |
| HRP-conjugated mouse anti-rabbit IgG | 1:10,000 | sc-2357, Santa Cruz Biotechnology | Figure 1b, 2h, 4h, 8e, 8g, 9h and supplementary figure 3b, 3e |
| DyLight 800 goat anti-Mouse IgG (H+L) | 1:10,000 | SA5-35521, ThermoFisher | Figure 9h |

a; Custom made by ProSci, Inc. using CDPQQSVVMRAVANLPLVSSTYDL, which is 7 to 28 amino acids of mouse Plin2 with addition of C at the beginning to increase the antigenicity.
HRP: horseradish peroxidase

**Supplementary Table 5: Donor ID of human islets used**

| Data | Source | | |
|---|---|---|---|
| | IIDP[a] | Alberta Institute Core[b] | Prodo[c] |
| Figure 8a-d | | | HP-17055<br>HP-17061<br>HP-17075 |
| Figure 8e Representative blot | SAMN08971735 | | |
| Figure 8e Densitometry | SAMN08971735 | | HP-17040<br>HP-17055<br>HP-17075 |
| Figure 8f, h | SAMN10737781<br>SAMN11476721<br>SAMN11483342<br>SAMN11864195 | | |
| Figure 8g Representative blot | SAMN10977276 | | |
| Figure 8g Densitometry | SAMN10977276<br>SAMN14120450<br>SAMN15400953<br>SAMN15579355 | SAMN12044342<br>SAMN15239415 | |
| Figure 8i, j | SAMN10737781<br>SAMN11476721<br>SAMN11483342<br>SAMN11633049<br>SAMN14132340<br>SAMN15400953<br>SAMN15579355 | SAMN15239415 | |
| Figure 9a | SAMN11476721 | | |
| Figure 9b-g | SAMN11476721<br>SAMN11483342<br>SAMN11633049<br>SAMN11864195<br>SAMN13938639<br>SAMN14120450<br>SAMN14132340<br>SAMN15400953<br>SAMN15579355 | | |
| Figure 9h Representative blot | SAMN15400953 | | |
| Figure 9h Densitometry | SAMN14120450<br>SAMN15400953<br>SAMN15579355 | SAMN15239415 | |

a; Integrated Islet Distribution Program (https://iidp.coh.org)
b: Alberta Diabetes Institute Islet Core (https://www.epicore.ualberta.ca/isletcore/)
c; Prodo laboratories INC (https://prodolabs.com)

**Supplementary Table 6: Donor and islet characteristics of human islets used**

| | Donor characteristics | | | | | Islet characteristics | | |
|---|---|---|---|---|---|---|---|---|
| ID | Age (years) | Sex (M/F) | BMI (kg/m$^2$) | HbA1c | Cause of death | Islet isolation center | Purity (%) | Viability (%) |
| SAMN 08971735 | 50 | M | 32.3 | 5.4 | Head trauma | The Scharp-Lacy Research Institute/IIDP | 90 | 95 |
| SAMN 10737781 | 66 | M | 27.2 | 4.7 | Cerebrovascular/stroke | The Scharp-Lacy Research Institute/IIDP | 95 | 95 |
| SAMN 10977276 | 52 | M | 27.2 | 5.7 | Cerebrovascular/stroke | Southern California Islet Cell Resource Center/IIDP | 85 | 96 |
| SAMN 11476721 | 50 | M | 32.8 | 6.0 | Anoxia | The Scharp-Lacy Research Institute/IIDP | 90 | 95 |
| SAMN 11483342 | 52 | F | 39.8 | 6.4 | Anoxia | University of Wisconsin/IIDP | 92 | 98 |
| SAMN 11633049 | 48 | M | 38.8 | 5.4 | Cerebrovascular/stroke | University of Wisconsin/IIDP | 90 | 98 |
| SAMN 11864195 | 53 | F | 40.0 | 5.5 | Cerebrovascular/stroke | University of Wisconsin/IIDP | 92 | 98 |
| SAMN 13938639 | 50 | F | 39.2 | 5.0 | Cerebrovascular/stroke | Southern California Islet Cell Resource Center/IIDP | 80 | 95 |
| SAMN 14120450 | 37 | M | 31.9 | 5.6 | Head trauma | University of Pennsylvania/IIDP | 85 | 96 |
| SAMN 14132340 | 31 | M | 27.0 | 5.2 | Head trauma | Southern California Islet Cell Resource Center/IIDP | 85 | 95 |
| SAMN 15400953 | 54 | F | 24.5 | 5.7 | Cerebrovascular/stroke | The Scharp-Lacy Research Institute/IIDP | 90 | 95 |
| SAMN 15579355 | 21 | M | 27.2 | 5.1 | Head trauma | The Scharp-Lacy Research Institute/IIDP | 85 | 95 |
| SAMN 12044342 | 44 | F | 23.2 | 4.9 | | Alberta Institute Core | | 90 |
| SAMN 15239415 | 31 | F | 20.3 | 4.8 | | Alberta Institute Core | | 95 |
| HP-17040 | 46 | M | 27.6 | 5.4 | Cerebrovascular/stroke | Prodo laboratories | 90 | 95 |
| HP-17055 | 41 | M | 28 | 5.3 | Head trauma | Prodo laboratories | 90 | 95 |
| HP-17061 | 38 | M | 39 | 5.8 | Anoxia | Prodo laboratories | | |
| HP-17075 | 37 | F | 19.8 | 4.6 | Anoxia | Prodo laboratories | 95 | 95 |
